# Volumetric Bioprinting of Bone-like Mineralizing Hydrogel Constructs in the Presence of High Cell Densities and Mineral Precursors

**DOI:** 10.1101/2025.07.03.662947

**Authors:** Bregje W.M. de Wildt, Margherita Bernero, Doris Zauchner, Ralph Müller, Xiao-Hua Qin

**Affiliations:** Institute for Biomechanics, ETH Zurich

**Keywords:** volumetric bioprinting, bioresin, biomineralization, *in vitro* model, bone, tissue engineering

## Abstract

A major challenge in bone tissue engineering is the embedding of osteocyte-like cells at high density within a mineralized matrix at the micro-scale and a trabecular-like architecture at the macro-scale. Volumetric bioprinting (VBP) enables rapid creation of complex cell-laden constructs through tomographic light projections. However, integrating both high cell densities and inorganic mineral precursors into VBP processes poses challenges due to light scattering, which can compromise print fidelity. In this study, we aim to combine bioinspired polymer-induced liquid-phase precursor (PILP) mineralization with VBP to fabricate cell-laden gelatin methacryloyl hydrogel constructs with amorphous mineral precursors. By stabilizing amorphous mineral precursors with poly-aspartic acid, light scattering is sufficiently reduced to enable printing. Tuning the refractive index of this mineralizing bioresin allows fast VBP of mineralized bone-like constructs with cell densities of up to 3 million cells ml^-1^. The constructs display high cell viability (>90%) and enhanced mineralization when cultured in osteogenic conditions with β-glycerophosphate. Encapsulated human mesenchymal stromal cells exhibit an early osteocytic phenotype after 28 days of differentiation. Collectively, this PILP-assisted VBP platform holds promise for the development of advanced *in vitro* bone models with more physiologically relevant architecture and cellular composition.

## 1. Introduction

Recent progress in bone bioengineering has led to the development of 3D tissue models that replicate the bone microenvironment *in vitro*.^[1–4]^ Despite these advances, difficulties remain in recapitulating native-like bone architecture, matrix composition, and cellular complexity.^[1]^ One major challenge is the embedding of osteocyte-like cells in a mineralized matrix (organic-inorganic composite) at the micro-scale and a trabecular-like architecture at the macro-scale.^[1,3,5]^ Current *in vitro* models often lack osteocytes, despite them being the most abundant cell type in bone, regulating remodeling through mechanotransduction and endocrine signaling.^[2,6,7]^ Recently, several advanced 3D *in vitro* models have been introduced in which osteocyte progenitor cells or cell lines have been seeded within or on top of collagen hydrogels,^[8–12]^ in between calcium phosphate microbeads and pillars,^[13–16]^ in porous silk fibroin scaffolds,^[17]^ or in spheroids.^[18–25]^ While some of these models do provide a bone-mimetic matrix allowing to study 3D cell-matrix interactions, they lack the embedding of cells within a trabecular-like architecture macroscopically.

Volumetric bioprinting (VBP) is an enabling technology to fabricate structurally complex cell-laden hydrogel constructs within tens of seconds through tomographic light projections.^[26]^ For instance, trabecular-like constructs have been fabricated with VBP using a gelatin methacryloyl (GelMA) or a photoclickable polyvinyl alcohol (PVA) as the bioresin.^[21,27]^ VBP was also used to fabricate thiol-ene gelatin constructs mimicking the osteoid niche^[22,28]^. Nevertheless, these constructs still require long-term culture to achieve mineralization. While post-printing mineralization methods show promise for biomimetic mineralization of cell-laden collagen and GelMA hydrogels,^[10,29,30]^ it could cause diffusion-dependent mineralization and cannot be applied for constructs in which mineralization is only desired locally such as for osteochondral and bone-tendon interfaces. A one-step biofabrication and mineralization approach is therefore needed.

The print fidelity in VBP depends on the ability to precisely control the spatial distribution of the effective light dose within the bioresin.^[31]^ The presence of scattering elements such as cells and mineral particles can significantly affect printability.^[31,32]^ As such, VBP of structurally complex mineralized hydrogels that also contain a high cell density remains an unsolved challenge.^[33]^ To address this, Bernal et al. have optically tuned GelMA bioresins using iodixanol to match the refractive indices of different (bio)resin components.^[31]^ In addition, Madrid-Wolff et al. have modified the light projections after characterizing the scattering within the resin.^[34]^ However, these methods only work with resins containing scattering elements with one defined refractive index (in case of refractive index matching) and a homogenous distribution of scattering elements in size and space (in case of correcting projections). Consequently, VBP with both high cell density and mineral particles remains difficult.

In this study, we set out to develop VBP of trabecular-like hydrogel constructs in the presence of high cell densities and mineral precursors. We employ the combination of polymer-induced liquid-phase precursor (PILP) mineralization and refractive index tuning to control scattering elements in a GelMA bioresin to enable VBP (**Figure 1**). *In vivo*, mineralization is regulated by acidic non-collagenous proteins (*e*.*g*., osteopontin, dentin matrix acidic phosphoprotein 1 (DMP1)), that control calcium and phosphate crystallization.^[35]^ *In vitro*, poly-aspartic acid (pAsp) mimics this role by stabilizing calcium and phosphate ions into amorphous calcium phosphate (ACP)^[36–38]^ via the PILP mechanism.^[36–40]^ We hypothesized that PILP mineralization could improve printability in the presence of calcium phosphate by reducing light scattering. Our results showed that pAsp was instrumental to control mineralization and consequently improve light transmission and printability. By combining PILP-mineralization with refractive index matching, we were able to fabricate trabecular-like constructs with an organic-inorganic composite matrix and a high cell density up to 3 million cells ml^-1^. Therefore, this methodology provides a foundation for the fabrication of biomimetic bone-like structures, paving the way for more physiologically relevant *in vitro* bone models.

**Figure 1.**
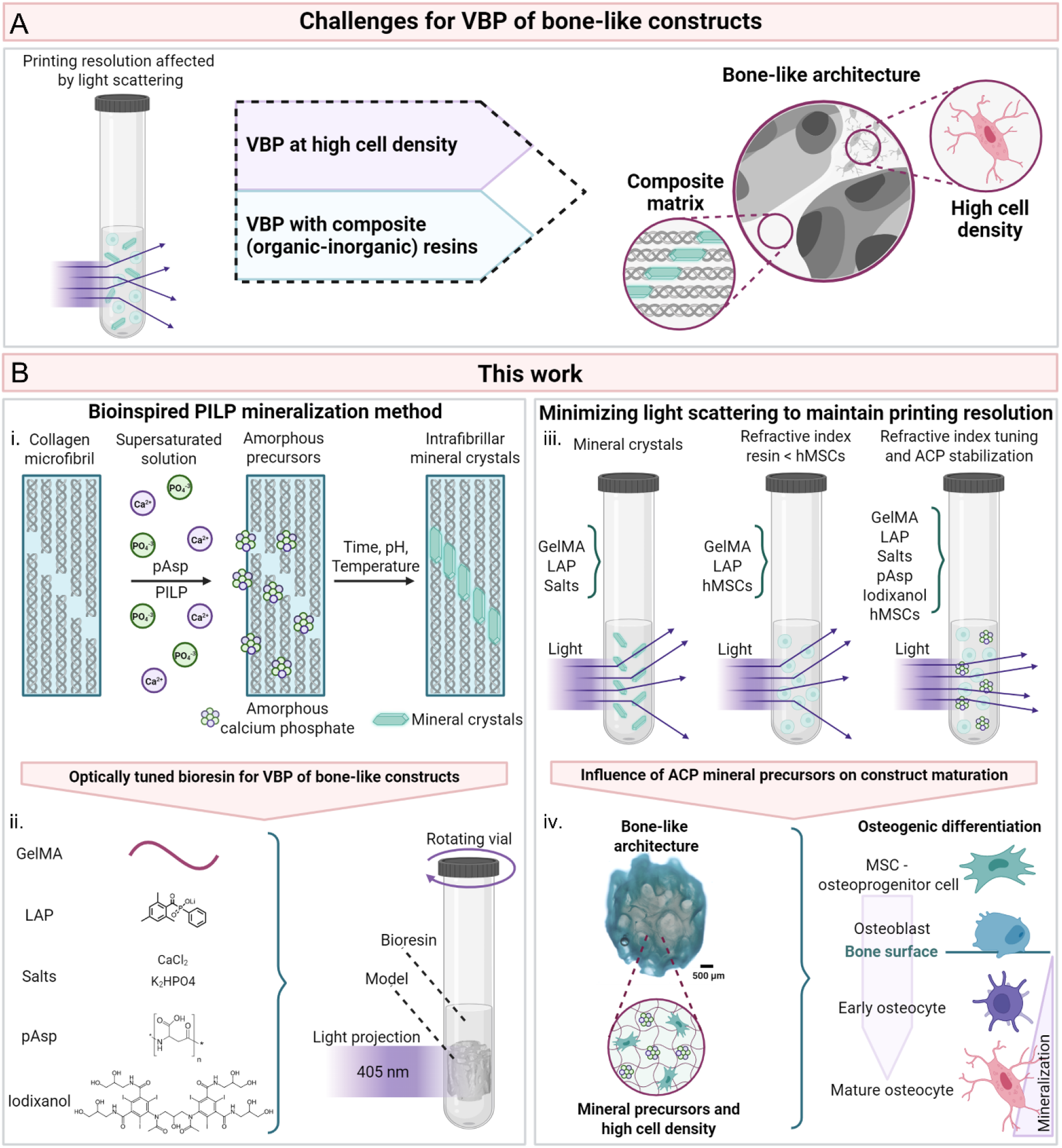
Rationale and study plan behind this work. (**A**) Challenges in VBP: VBP allows for fast photofabrication of trabecular-like architectures, but bioprinting in the presence of light-scattering elements, such as mineral particles and high cell densities, remains challenging. (**B**) Illustration of the methodologies in this work. We leverage polymer-induced liquid precursor (PILP) biomineralization to stabilize ACP (i). By converging the PILP method with optical refractive index tuning (ii), we sought to reduce light scattering to enable VBP of constructs with trabecular bone-like architecture, high cell density, and a mineralized matrix (iii). These constructs were subsequently evaluated for their ability to promote osteocytic differentiation and matrix mineralization (iv). Figure created with BioRender.com. Abbreviations: volumetric bioprinting (VBP), polymer-induced liquid precursor phase (PILP), gelatin methacryloyl (GelMA), Lithium-Phenyl-2,4,6-trimethylbenzoylphosphinat (LAP), poly-aspartic acid (pAsp), human mesenchymal stromal cells (hMSCs), amorphous calcium phosphate (ACP).

## 2. Results and discussion

### 2.1 Stabilization of amorphous mineral precursors in GelMA bioresins with PILP

The bioresin consists of 5% GelMA, 0.1% lithium-phenyl-2,4,6-trimethylbenzoylphosphinat (LAP) photoinitiator and pAsp for PILP-induced mineralization. GelMA was used as the hydrogel precursor due to its excellent biocompatibility and suitability for VBP.^[27]^ A 5% GelMA concentration was chosen to allow permissive culture conditions for cell spreading – an important hallmark of osteocytic differentiation.^[22]^

First, the required concentration of pAsp was determined by measuring the resins’ optical densities (OD) at the wavelength (*λ* = 405 nm) of VBP. Varying concentrations of CaCl_2_ (low L: 9 mM, medium M: 18 mM, high H: 27 mM), K_2_HPO_4_ (low L: 4.2 mM, medium M: 8.4 mM, high H: 12.6 mM) and pAsp (low L: 100 µg ml^-1^or high H: 1 mg ml^-1^.) were included (**Table S1**). As PILP mineralization is optimal at physiological pH and temperature, the pH was controlled by preparing the resins with HEPES (25 mM) as buffering salts at pH 7.4, and resins were incubated at 37 °C before further use. After 1h incubation at 37 °C, resins were pipetted in clear 96-well plates and cooled to 4 °C (eventual printing temperature) before measuring the OD. While the addition of pAsp alone did not affect the resin’s final OD, 1 mg ml^-1^pAsp could significantly reduce the OD at all salt concentrations (**Figure S1**), which could be attributed to reduced light scattering or absorption by the resin. Therefore, we chose to use 1 mg ml^-1^pAsp in the following experiments.

As time can also influence mineral nucleation and further crystallization, we characterized the time-dependent OD at 37 °C for the conditions with and without 1 mg ml^-1^pAsp (**Figure 2A**). LAP was omitted to prevent unwanted premature crosslinking. The formation of calcium phosphate clusters likely increased the resins’ OD over time, which happened faster when higher salt concentrations were used in the absence of pAsp (**Figure 2B**). This phenomenon might be explained by the classical and non-classical nucleation theory. In the resins with only salts, calcium and phosphate must overcome an ion concentration dependent energy barrier before they nucleate. After forming a stable nucleation cluster, mineral crystals grow rapidly, likely until the resin is no longer supersaturated resulting again in a stable OD after the rapid increase.^[41]^ The addition of pAsp resulted in a largely flattened OD for all salt concentrations (**Figure 2B**). After 30 min of incubation, all salt concentrations reached a relatively stable OD. As such, when pAsp is added to the resins, it is expected that calcium and phosphate rapidly form an intermediate mineral precursor that remains stable over 1 h of incubation, providing a sufficient time-window for VBP. The 30 min incubation time was used in the following experiments.

**Figure 2.**
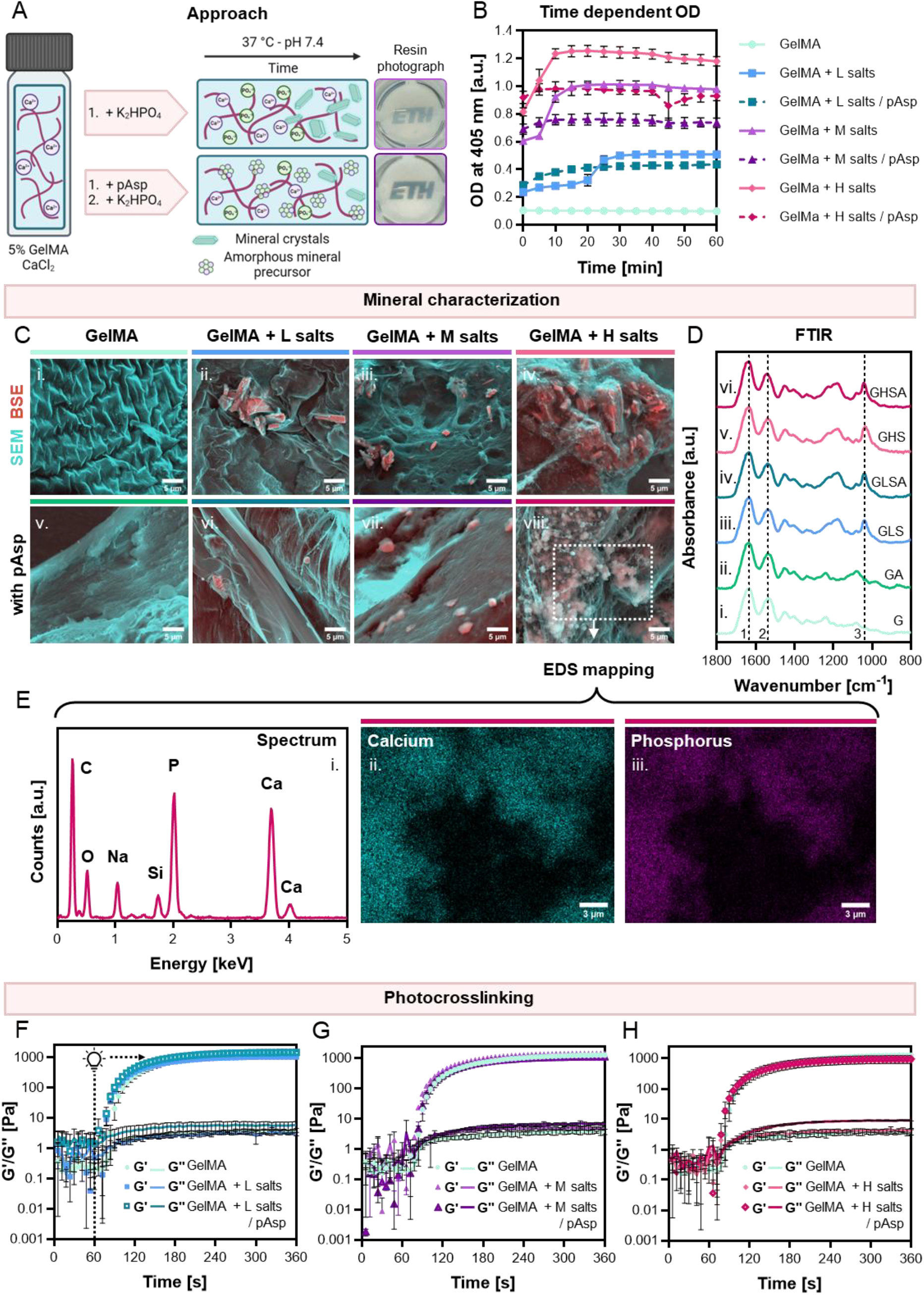
Characterization of resins doped with mineral precursors. (**A**) By adding pAsp and inorganic salts to GelMA, small ACPs are stabilized in the GelMA resins to enable light transmission while maintaining printing resolution. Figure created with BioRender.com. (**B**) Time-dependent OD of different GelMA resins as an influence of salt concentration and pAsp. Data shown as mean ± standard deviation, *N* = 4. (**C**) Mineral morphology visualized with SEM with BSE detection demonstrating plate-like mineral crystals in the absence of pAsp (ii - iv) and amorphous-like minerals in the presence of pAsp (vi - viii), *N* = 3. (**D**) representative FTIR spectra of GelMA (i - G), GelMA + pAsp (ii - GA), GelMA + low salt concentration (iii - GLS), GelMA + low salt concentration + pAsp (iv - GLSA), and high salt concentrations without (v - GHS) and with pAsp (vi - GHSA), with evidence for amide I (1), amide II (2), and phosphate stretching vibration (3) peaks, *N* = 3. (**E**) EDS mapping of area indicated in C (viii) with evidence for calcium (ii) and phosphorous (iii) within the amorphous-like mineral particles. (**F-H**) Time sweep of photorheology: storage and loss moduli measured under UV-light (λ = 365 nm) from 60 s onwards of resins with low, medium, and high salt concentrations, respectively. Data represented as mean ± standard deviation, *N* = 3. Abbreviations: poly-aspartic acid (pAsp), gelatin methacryloyl (GelMA), optical density (OD), low (L), medium (M), high (H), scanning electron microscopy (SEM), backscattered electron (BSE), Fourier-transform infrared spectroscopy (FTIR), energy dispersive spectroscopy (EDS).

To evaluate hydrogel mineralization, 5% GelMA hydrogels with and without salts and 1 mg ml^-1^pAsp were cast by *in situ* photocrosslinking. The impact of salt concentration was investigated in terms of 1) mineral morphology using scanning electron microscopy (SEM) with backscattered electron (BSE) detection, 2) chemical composition using Fourier-transform infrared spectroscopy (FTIR), and 3) Ca and P content using energy dispersive spectroscopy (EDS). When visualizing mineralization, the GelMA only and GelMA + pAsp compositions exhibited no signs of mineralization as indicated by morphology and limited BSE signal (**Figure 2C**). The backscattering of electrons happens more in denser structures and features with higher atomic mass such as Ca and P in comparison to the C, O and N in GelMA. Indeed, in resin compositions with salts but without pAsp, plate-like mineral crystals were observed with high BSE signal in a dose-dependent manner. In resin compositions with both salts and pAsp a similar dose-dependent response was observed. However, minerals appeared more amorphous-like with less intense BSE signal in comparison to the compositions without pAsp (**Figure 2C**). FTIR spectra of all compositions confirmed the presence of GelMA-corresponding amide peaks at ∼1635 cm^-1^(amide I) and ∼1540 cm^-1^(amide II) (**Figure 2D, Figure S2**). Additionally, in all salt-containing compositions, a [PO_4_]^3-^ stretching vibration peak at ∼1040 cm^-1^ indicated the presence of phosphate. The presence of Ca and P was confirmed in all salt-containing compositions as observed in the EDS spectral area analyses (**Figure 2E, Figure S3**).

Next, we characterized the influence of salts/pAsp inclusion on the photocrosslinking and mechanical properties of GelMA hydrogels. By measuring and analyzing the storage (G’) and loss (G’’) moduli under UV irradiation over a period of 5 min, hydrogel formation was observed in all compositions **(Figure 2F-H, Figure S4)**. However, no clear trends were observed in the G’ plateau likely due to the change of refractive indices in the resins (**Figure S5**). The addition of pAsp and salts led to a reduction in time required to reach 85% of the G’-plateau. The presence of small mineral particles might add an influence on the crosslinking. Next, we evaluated the mechanical and swelling properties of GelMA hydrogels. The addition of salts and pAsp led to a salt dose-dependent decrease in hydrogel stiffness measured with a compression test (**Figure S5**). The results show that the inclusion of mineral particles results in the decrease of gel stiffness. The mass swelling ratio was not altered by the addition of salts and pAsp. The GelMA + pAsp (no salts) group exhibited the highest degree of swelling.

To evaluate the cytocompatibility of the resins, human mesenchymal stromal cells (hMSCs) were photoencapsulated in the resins at a concentration of 3 million cells ml^-1^using photocrosslinking. After 48 hours of culture, high cell viability (>90%) was observed in the pure GelMA, GelMA + pAsp, and GelMA + low salts ± pAsp compositions (**Figure S6**). At higher salt concentrations, pAsp seems to add a protective effect on cell viability, as reported elsewhere.^[10]^ During a 21 day osteogenic culture, the addition of low concentration of salts and pAsp (GelMA + ACP) did not affect cell viability, as evidenced by a MTS assay (**Figure S7A-B**). Compared to the GelMA group, cells exhibit a similar increase in metabolic activity from day 3 to day 21. No changes in DNA content and lactate dehydrogenase (LDH) release were observed in the culture medium (**Figure S7C-D**). However, clear differences were observed when evaluating calcium content and alkaline phosphatase (ALP) activity (**Figure S7E-F**). After 21 days of culture, cells in the GelMA + ACP group showed significantly higher calcium content and ALP activity than those in the GelMA control group (**Figure S7G**).

### 2.2 The influence of resin mineralization on printability

It was expected that the stabilization of ACP with pAsp could reduce light scattering during VBP (**Figure 3A**). To study the light-resin interactions, we first quantified light transmission (405 nm) through the resin over a period of 30 s using a power meter (**Figure 3B**). LAP was omitted to prevent possible optical changes as a result of photocrosslinking. The presence of salts in the resins resulted in significant lower light transmission, which could be prevented by the addition of pAsp (**Figure 3C**). In conditions with medium and high salt concentrations, almost no light was transmitted through the resins (**Figure 3B-C**), which was also indicated by visual inspection of the cuvettes filled with resins (**Figure 3D**). By projecting a spatially coherent light beam on the vials and by taking photographs with the orthogonal camera within the printer, we visualized resin-induced light scattering.^[34]^ Indeed, in resins with salts, light was scattered to varying degrees as an influence of the compositions (**Figure 3D**). Light beams diverged less in the presence of pAsp in comparison to their control groups. As light transmission was poor in resins with medium and high salt concentrations, only the low salt concentration group was used for further experiments.

**Figure 3.**
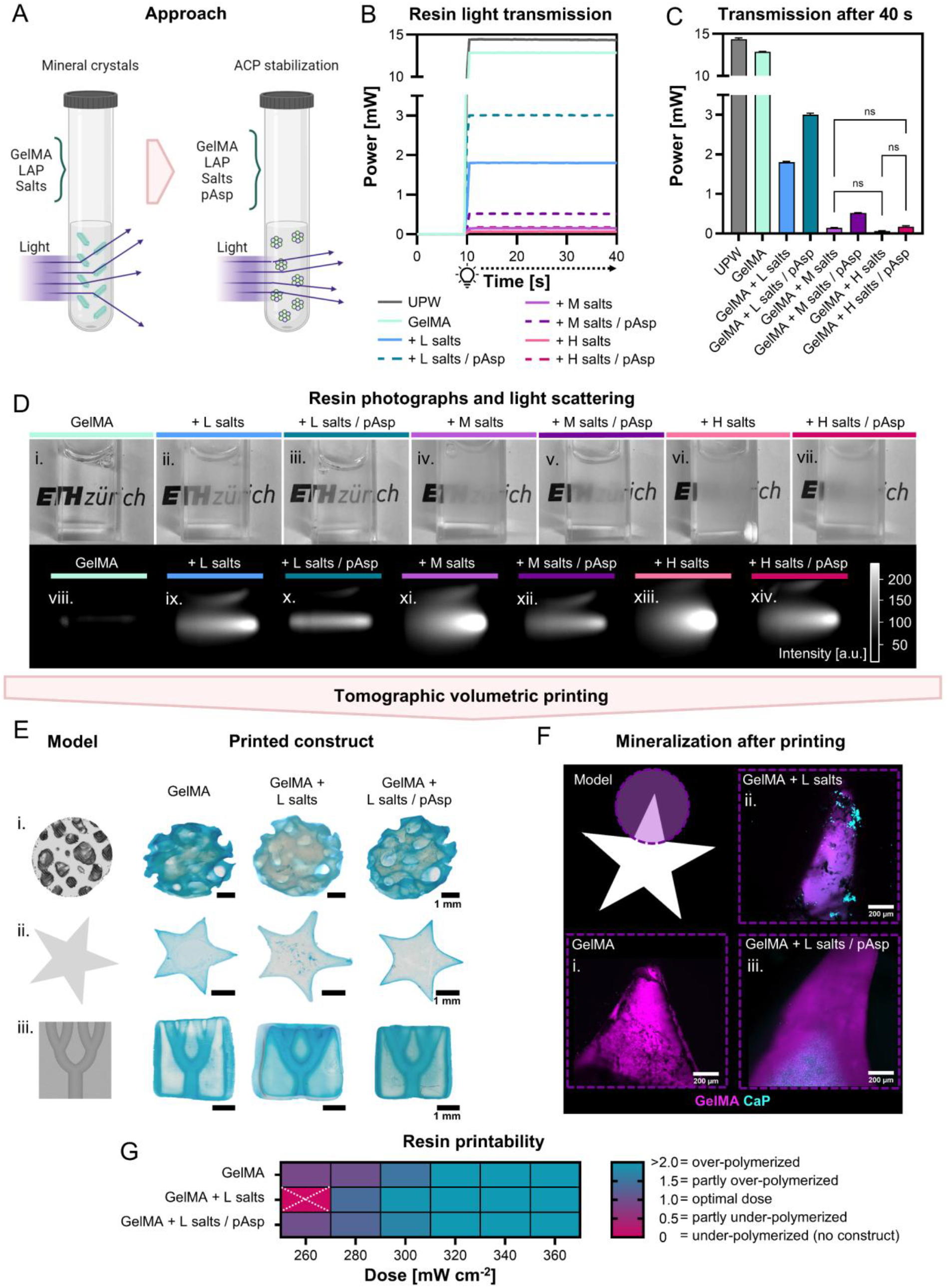
Influence of mineralizing resins on volumetric printing. (**A**) Photo-curable mineralizing GelMA resins were prepared and characterized for their light transmission properties and printability. Figure created with BioRender.com. (**B**) Light transmission through resins measured over 30 s (typical printing duration) within the printer. Data shown as mean values, *N = 3*. (**C**) Light transmission through resins at 40 s, mean ± standard deviation, *N* = 3, *p*<0.05 (One-way ANOVA and Holm-Šídák’s post hoc tests, only non-significant differences presented. All other pairs were statistically significantly different). (**D**) Resin photographs (i - vii) and scattered light within the resins visualized by projecting a spatially coherent light beam on the resin-filled vials and by photographing with an orthogonal camera within the printer (viii - xiv), *N* = 3. (**E**) Tomographic volumetric printing of a trabecular bone model (i), star (ii), and branch (iii), *N* ≥ 3. (**F**) Confocal microscopic images of rhodamine -labelled GelMA and Osteoimage-stained calcium phosphate, *N* = 3. (**G**) Quantification of dimension ratios between the branch model branch diameter in the model and the actual printed diameter as an influence of printing doses, *N* = 3. Abbreviations: gelatin methacryloyl (GelMA), poly-aspartic acid (pAsp), Lithium-Phenyl-2,4,6-trimethylbenzoylphosphinat (LAP), ultrapure water (UPW), amorphous calcium phosphate (ACP), low (L), medium (M), high (H), not significant (*ns*).

Next, we studied the influence of low salt concentration and pAsp on volumetric printing of different models, including trabecular bone, a star, and a branch model. **Figure 3E** shows the constructs with improved printing fidelity in the presence of pAsp. In contrast, the constructs without pAsp exhibited both over- and under-polymerized features. To quantify the printability, we fabricated the branch model with each resin at 6 different doses, measured the diameter of the central channel, and determined the ratio between the model diameter and the printer construct diameter (**Figure 3G**). A ratio of 1 suggests an optimal dose, whereas a ratio above or below 1 indicates over-polymerization or under-polymerization, respectively. Constructs with only salts resulted in either over-polymerization or under-polymerization, while constructs with both salts and pAsp showed a similar dose-response as GelMA (**Figure 3G**). When printing at 260 mW cm^−2^, no construct was accessible for the GelMA + salts group, indicating compromised printability (**Figure S8**).

Since the size and distribution of ACP are key to hydrogel mineralization, we further performed confocal imaging of rhodamine-labelled GelMA matrix and hydroxyapatite minerals (CaP). For the GelMA + salts group, large mineral aggregates were observed (**Figure 3F**). The addition of pAsp showed much smaller but more homogeneously distributed minerals. However, less mineralization was observed at the edges of the constructs, indicating the diffusion of mineral precursors. These findings imply that the printed GelMA matrices may provide a permissive environment to support the nucleation and maturation of ACP clusters into hydroxyapatite. In collagen, mineral precursors are believed to get “entrapped” within gap regions in between the triple helices. It has been hypothesized that in the absence of pAsp, extrafibrillar mineralization has the lowest nucleation barrier and is therefore preferred over intrafibrillar mineralization through the collagen gap regions.^[42]^ In the presence of pAsp the nucleation barrier in extrafibrillar spaces increases while the gap region provides a confined space that reduces the nucleation barrier, favoring nucleation inside the gap region.^[42,43]^ In addition to the role of confinement in nucleation, positively charged regions within the collagen gap zone have been suggested to attract negatively charged pAsp-ACP clusters.^[44]^ Within our GelMA hydrogels, mineral stabilization and nucleation could thus either occur through the hydrogel mesh, providing locally confined spaces upon polymerization, or by providing positively charged (non-methacrylated lysine residues) regions that bind pAsp-ACP complexes.^[30,44,45]^ Given the differences between photocrosslinked GelMA and natural collagen, future work is warranted to dissect the exact PILP-induced mineralization mechanisms in printed GelMA matrices. As volumetric printing was difficult in the absence of pAsp, we continued our study using GelMA + salts + pAsp and GelMA (control).

### 2.3 Optically tuned VBP of mineralizing hydrogels at high cell density

In addition to mineral particles, the presence of cells at high densities can also significantly affect print fidelity during VBP. To enable VBP of both high cell densities and mineral precursors, we combined our PILP mineralization with optical tuning as reported by Bernal et al (**Figure 4A**).^[31]^ In this approach, iodixanol is used as a sacrificial additive to increase the resin’s refractive index to a level that matches the refractive index of cells.^[31]^ Cell organelles and intracellular matter have various refractive indices ranging from 1.355 – 1.600.^[46]^ Aimed at this range, we mixed GelMA and GelMA + ACP with 10%, 15%, and 20% iodixanol and measured the resin’s refractive indices. Each concentration of iodixanol can increase the refractive index of resin groups GelMA and GelMA + ACP (**Figure 4B**). At 10%, the resins’ refractive indices reached the range of cellular components (**Figure 4B**). To evaluate the influence of this increase in refractive index on light transmission, optically tuned resins were mixed with 5 million cells ml^-1^. The resulting OD was measured at 405 nm using a plate reader. For the GelMA group, the OD decreased with all iodixanol concentrations, but no statistical differences between the groups (**Figure 4C**). In the GelMA + ACP group, the OD remained the same with the addition of iodixanol, despite the increase in refractive index (**Figure 4B-C**). However, when visualizing light scattering within the bioresins (3 million cells ml^-1^), the addition of 10% iodixanol reduced the light scattering in both resin types (**Figure 4D**).

**Figure 4.**
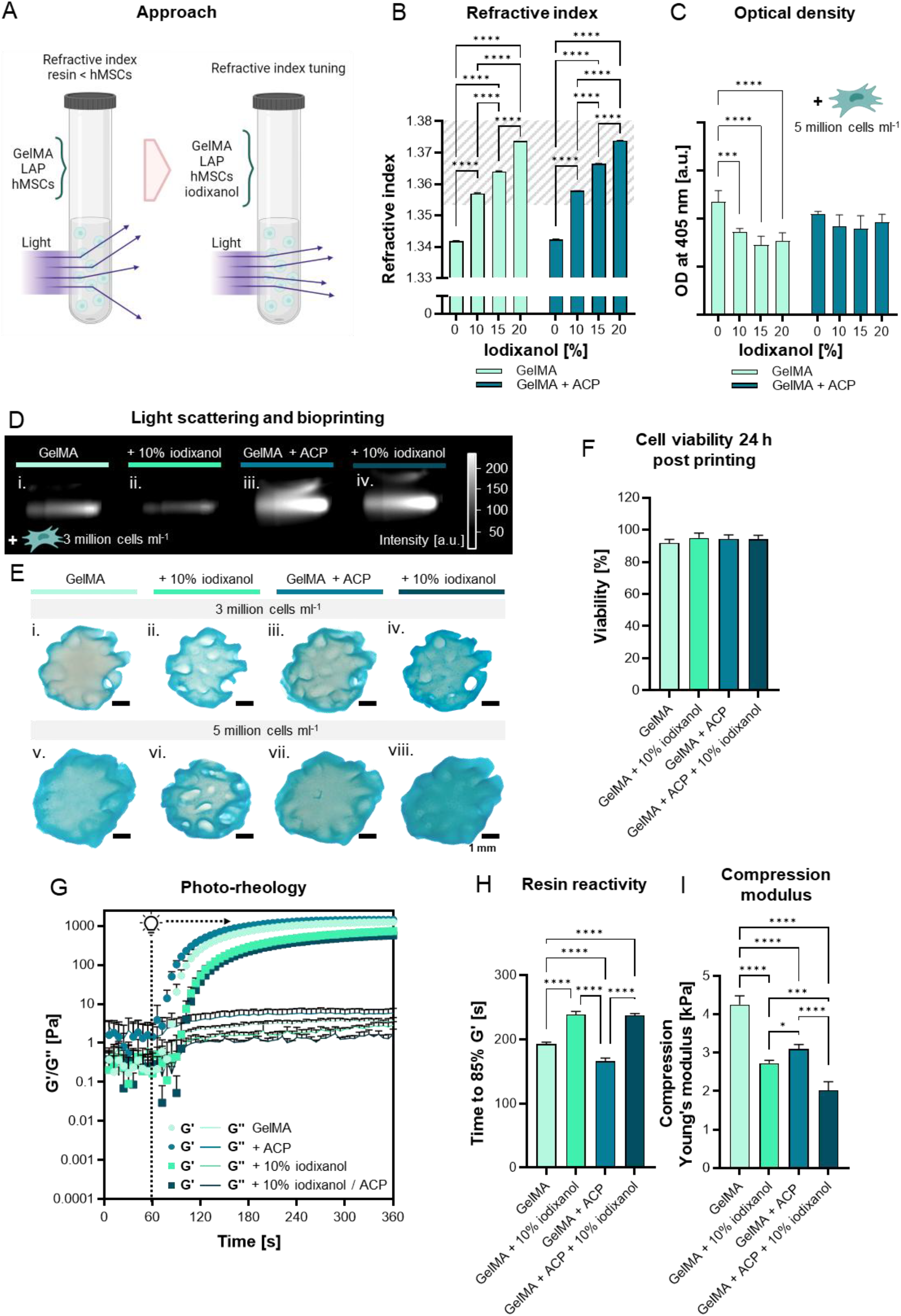
Combining PILP mineralization with refractive index tuning for VBP at high cell density. (**A**) Matching the refractive index of pure GelMA and GelMA + ACP resins with the refractive index of hMSCs using iodixanol to improve printability. Figure created with BioRender.com. (**B**) Influence of iodixanol on the refractive indices of GelMA and GelMA + ACP resins. The grey marked area refers to the known range for cellular components. Data shown as mean ± standard deviation, *N* = 3, *p*<0.05 (Two-way ANOVA and Tukey’s post hoc tests for comparisons between iodixanol concentrations within pure GelMA and GelMA + ACP resins). (**C**) Resins’ OD in the presence of 5 million cells ml^-1^and iodixanol, mean ± standard deviation, *N* = 4, *p*<0.05 (Two-way ANOVA and Tukey’s post hoc tests for comparisons between iodixanol concentrations within pure GelMA and GelMA + ACP resins). (**D**) Scattered light within the resins in the presence of 3 million cells ml^-1^, *N* = 3. (**E**) Tomographic volumetric printing of a trabecular bone model at a cell density of 3 million cells ml^-1^(i - iv) and 5 million cells ml^-1^(v - viii), *N* ≥ 2. (**F**) Cell viability quantification of hMSCs (3 million cells ml^-1^) following 24 h after VBP. Data shown as mean ± standard deviation, *ns* (One-way ANOVA and Holm-Šídák’s post hoc tests). (**G**) Photorheology of different resins showing the evolution of storage and loss moduli measured under influence of UV-light (λ = 365 nm) from 60 s onwards, mean ± standard deviation, *N* = 3. (**H**) Photo-rheology derived time required to reach 85% of G’-plateau storage modulus, mean ± standard deviation, *N* = 3, *p*<0.05 (One-way ANOVA and Holm-Šídák’s post hoc tests). (**I**) Young’s modulus derived from compression stress-strain curves, mean ± standard deviation, *N* ≥ 3, *p*<0.05 (One-way ANOVA and Holm-Šídák’s post hoc tests). Asterisks in figures represent results of post hoc analyses (\**p*<0.05, \*\**p*<0.01, \*\*\**p*<0.001, \*\*\*\**p*<0.0001). Abbreviations: gelatin methacryloyl (GelMA), Lithium-Phenyl-2,4,6-trimethylbenzoylphosphinat (LAP), amorphous calcium phosphate (ACP), optical density (OD), human mesenchymal stromal cells (hMSCs), not significant (ns).

Next, we evaluated the influence of iodixanol on the photocrosslinking and mechanical properties of GelMA hydrogels by photorheology (**Figure 4G**). Photocrosslinking of GelMA with 10% iodixanol resulted in significantly lower G’-plateau value for both groups (**Figure S9**), as reported elsewhere.^[31,47]^ Moreover, the addition of 10% iodixanol led to an increase in time required to reach 85% of the G’-plateau, suggesting slower photocrosslinking in the presence of iodixanol (**Figure 4H**). In line with the decrease in G’-plateau, the addition of iodixanol also led to a decrease in hydrogel stiffness measured with a compression test, with the lowest stiffness for GelMA hydrogels with ACP and iodixanol (**Figure 4I**).

When evaluating resin printability at 3 million cells ml^-1^and 5 million cells ml^-1^, resins with iodixanol led to better printability, especially for pure GelMA compositions at both cell densities (**Figure 4E**). In trabecular constructs printed with 10% iodixanol, pore architectures could be resolved, whereas no pores were visible in the pure GelMA without iodixanol. For GelMA + ACP resins, iodixanol improved printability to a lesser extent and only for 3 million cells ml^-1^(**Figure 4E**). At 5 million cells ml^-1^, bioprinting with GelMA + ACP resins led to poorly defined constructs. Therefore, the following experiments were performed at the lower cell density of 3 million cells ml^-1^. Since prints with 10% iodixanol yielded very soft constructs, we performed post-curing to improve construct structural integrity in further experiments.

To verify if post-curing may induce cellular oxidative stress, we labeled hMSCs with 2′,7′-Dichlorodihydrofluorescein diacetate (DCFH-DA), which could be converted to fluorescent DCF in viable cells after oxidation by free radicals or reactive oxygen species.^[48]^ Labeled hMSCs were mixed into GelMA and GelMA + ACP resins in the presence and absence of iodixanol. Quantification of DCF-positive cells revealed a trend towards an increase in oxidative stress as a result of iodixanol (**Figure S10**). In most conditions, post-curing did not change the number of DCF-positive cells. Therefore, post-curing was a useful step to stabilize constructs when resins were combined with iodixanol. A subsequent cell viability assay at 24 h confirmed >90% viability in all post-cured constructs (**Figure 4F, Figure S11**).

### 2.4 Influence of the PILP resin formulation on osteogenic differentiation of hMSCs

Composite hydrogels have been shown to promote osteogenic differentiation of hMSCs.^[10,37]^ To investigate whether calcium phosphate may have an impact on osteogenic differentiation, we cultured bioprinted constructs for up to 28 days (**Figure 5A**). GelMA and GelMA + ACP constructs with 3 million cells ml^-1^ were successfully bioprinted in the presence of 10% iodixanol and post-cured, leading to trabecular constructs with resolved pore structures (**Figure S12**). Cells remained highly viable for 28 days within all bioprinted constructs. This was further evidenced by low LDH activity in the medium (**Figure 5B**), negligible change in DNA content (**Figure S13**), and an increase in metabolic activity towards day 28 (**Figure 5C**).

**Figure 5.**
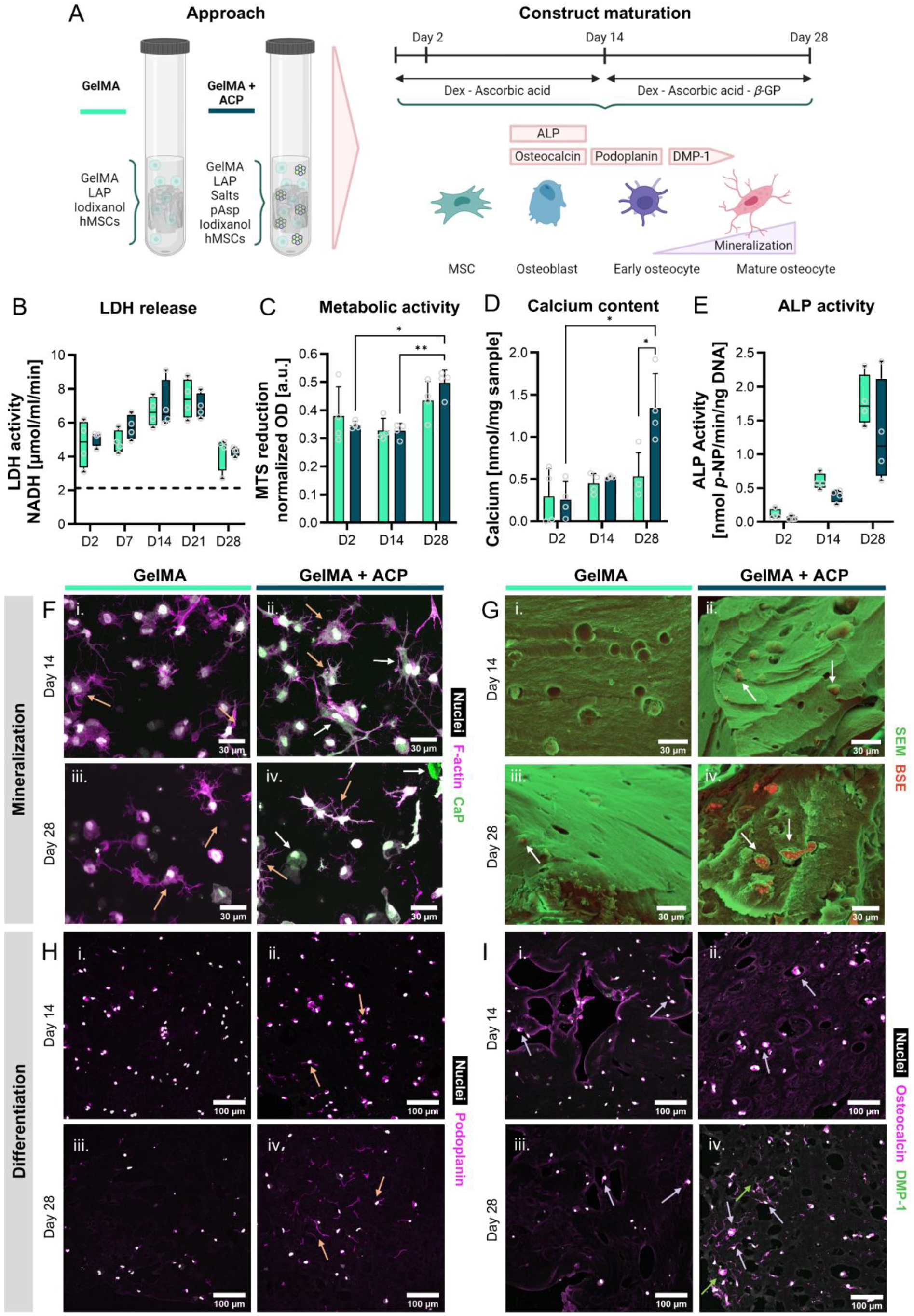
Influence of the mineralizing resin on osteogenic differentiation of hMSCs after VBP. (**A**) Constructs were bioprinted and cultured for up to 28 days with first 14 days osteogenic medium without beta-GP followed by 14 days with beta-GP. Figure created with BioRender.com. (**B**) LDH release in the medium as a measure for cell death. Data shown as median ± interquartile range, *N* = 4, *ns* (Mann-Whitney U tests per time point with Bonferroni correction for multiple comparisons). (**C**) Cell metabolic activity measured by MTS reduction, mean ± standard deviation, *N* = 4, *p*<0.05 (Two-way ANOVA and Tukey’s post hoc tests). (**D**) Construct mineralization measured by calcium content, mean ± standard deviation, *N* = 4, *p*<0.05 (Two-way ANOVA and Tukey’s post hoc tests). (**E**) Cellular ALP activity over time. Data shown as median ± interquartile range, *N* = 4, *ns* (Mann-Whitney U tests per time point with Bonferroni correction for multiple comparisons). (**F**) Confocal images of cell spreading and matrix mineralization after staining for cell nuclei, F-actin, and hydroxyapatite, *N* = 3. (**G**) Microscopic images of cells and minerals visualized with SEM with BSE detection, *N* = 3. (**H**) Confocal images of stained cells after staining for nuclei and podoplanin in thick tissue sections, *N* = 3. (**I**) Confocal images of osteocalcin and DMP1 markers in cells, *N* = 3. Asterisks in figures represent results of post hoc analyses (\**p*<0.05, \*\**p*<0.01). Abbreviations: dentin matrix acidic phosphoprotein 1 (DMP1), lactate dehydrogenase (LDH), alkaline phosphatase (ALP), calcium phosphate (CaP), and β-Glycerophosphate (beta-GP).

As β-glycerophosphate (beta-GP) may interfere with mineralization of GelMA hydrogels (**Figure S7**), we chose to omit beta-GP when evaluating the osteogenic potential of our bioprinted constructs for 14 days. ^[3,49,50][51,52]^This facilitates the evaluation of the influence of ACP on GelMA mineralization. After 14 days, beta-GP was included in the osteogenic medium (**Figure 5A**) to promote osteoblast to osteocyte transition.^[53]^ Interestingly, after 2 and 14 days of culture, no differences in calcium content were observed between the two hydrogel compositions (**Figure 5D**), which was also observed for cast hydrogels (**Figure S7**). These results imply that the mineral particles may diffuse out from the constructs due to their small sizes (**Figure 3F**). Nonetheless, when beta-GP was added to the medium, GelMA + ACP constructs were still able to accumulate significantly more calcium than GelMA alone constructs (**Figure 5D**), indicating a synergistic effect of the mineral precursors in the construct and beta-GP in the medium on mineralization. Both hydrogel compositions led to an increase in ALP activity over time, with no differences between the two resin types (**Figure 5E, Figure S13**).

By staining for hMSCs and CaP, we observed cell morphogenesis in both hydrogel compositions from day 14 (**Figure 5F**). Cell spreading and dendrite formation is a hallmark of osteocytic differentiation.^[54]^ Cell spreading was observed in the bioprinted constructs. Interestingly, in GelMA + ACP constructs, CaP appeared extracellularly in the matrix on days 2, 14 and 28, as well as within the cells on days 14 and 28 (**Figure 5F, Figure S14**). This indicates cellular phagocytosis of the minerals present in the hydrogels.^[61,62]^ Another explanation might be the intracellular formation of matrix vesicles after calcium and phosphate uptake from the environment.^[63,64]^ However, non-reacted calcium and phosphate are expected to wash out after printing. With SEM, we identified cells within lacunae (**Figure 5G**). The BSE signal confirmed the presence of cellular mineral particles mainly in GelMA + ACP constructs.

Next, we evaluated the influence of mineral-containing hydrogels on hMSC osteogenic differentiation using immunofluorescent staining. After 14 days of culture, cells in the GelMA + ACP constructs exhibit more podoplanin (early osteocyte marker) than in the GelMA constructs (**Figure 5H**). *In vivo*, early osteocytes form dendrites upon embedding, which is dependent on podoplanin expression.^[54,65,66]^ Indeed, podoplanin was found within the cell dendrites (**Figure 5H, Figure S15**). Osteocalcin has both been found in osteoblasts and osteocytes.^[66]^ In our constructs we found osteocalcin mainly in the cell bodies and dendrites of GelMA + ACP constructs at both time points (**Figure 5I, Figure S16**). Another osteocytic marker (DMP1) was also seen in cell bodies and dendrites in GelMA + ACP constructs on day 28 (**Figure 5I, Figure S17**). This indicates that the inorganic-organic matrix composition may have facilitated the osteocytic differentiation of hMSCs. Nevertheless, the slightly lower stiffness in GelMA + ACP constructs may also contribute to this difference (**Figure 4I, Figure S5**). In the future, these mineralizing cell-laden constructs may be combined with mechanical loading to further improve mineralization and osteocytic differentiation.^[23]^

## 3. Conclusion and Future Outlook

Taken together, we developed a methodology to print trabecular bone-like living constructs by VBP in the presence of high cell densities and mineral precursors. We leveraged the advantages of small sized ACP and optical tuning to reduce light scattering during VBP and induce hydrogel mineralization through PILP. The constructs containing mineral precursors demonstrate high cell viability and more mineralization when beta-GP was used in the culture medium. Within these constructs, hMSCs showed key signature markers corresponding to early osteocytic differentiation following differentiation for up to 28 days. Therefore, this PILP-assisted 3D bioprinting technique opens new avenues for future near-physiological *in vitro* models of human bone, where osteocyte-like cells are embedded in a mineralized matrix across a trabecular-like architecture. The convergence with other methods, such as phase separation,^[55,56]^ microgels,^[57]^ void templating,^[21,58]^ or two-photon ablation^[47,59,60]^ may eventually enable researchers to reconstruct the osteocyte lacuno-canalicular network found in natural bone in future.

## 4. Experimental section

### 4.1 Materials

GelMA (degree of substitution ≈58%) was synthesized by reacting gelatin (Type A, porcine skin; G2500, Sigma Aldrich, Schnelldorf, Germany) with methacrylic anhydride (276685, Sigma Aldrich) as previously described,^[67]^ dialyzed against UPW for 5 days, sterile filtered, lyophilized and stored at -20 °C until further use. Before experiments, a 25% w/v GelMA (5x) solution was freshly prepared in UPW. For photoinitiated GelMA crosslinking, LAP (900889, Sigma-Aldrich) was used, and a stock solution was prepared by dissolving LAP in UPW at a 0.5% w/v (5x) concentration. For post-curing, LAP was prepared in Dulbecco’s phosphate-buffered saline (D-PBS; 14040133, Thermo Fisher Scientific, Reinach, Switzerland) at a 1% (10x) concentration. Stock salt solutions (10x) were prepared by dissolving CaCl_2_ * 2H_2_O (C7902, Sigma-Aldrich) at concentrations of 90 mM, 180 mM and 270 mM, and K_2_HPO_4_ (P8281, Sigma-Aldrich) at concentrations of 42 mM, 84 mM and 126 mM in 250 mM HEPES (15630080, Thermo Fisher Scientific) in UPW. After dissolving, the pH of the salt stock solutions was set to 7.4 using 1 M HCl and NaOH. The solutions were stored at 4 °C for up to one month. A 100 mg/ml stock solution of pAsp (P3418, Sigma-Aldrich) was prepared (100x) in UPW, aliquoted and stored at -20 °C until further use. A 5x PBS (18912014, Thermo Fisher Scientific) solution was prepared and used for the non-mineralizing resins. For refractive index tuning, different concentrations (0%, 10%, 15% and 20% w/v) of iodixanol (OptiPrep^TM^; 07820, StemCell Technologies, Cologne, Germany) were used. For sterile resin preparation, LAP, salt and pAsp solutions were sterile filtered.

### 4.2 Resin preparation

Control resins (5% GelMA and 0.1% LAP in 1x PBS) were prepared by mixing UPW 2:5, 5x PBS 1:5, 25% GelMA 1:5 with 0.5% LAP 1:5. Mineralizing resins (5% GelMA, salts with or without pAsp and 0.1% LAP in UPW) were prepared by mixing in the following order UPW (with or without pAsp 1:100) 2:5, CaCl_2_ stock solution 1:10, 25% GelMA 1:5 with 0.5% LAP 1:5 and K_2_HPO_4_ stock solution 1:10. Before VBP, resins were incubated for 30 min at 37 °C in air-tight tubes protected from light to allow for resin mineralization. For iodixanol containing resins, iodixanol was mixed into the UPW – pAsp - CaCl_2_ premix (**Table S1**). For cellular constructs, resins were prepared sterile and after mineralization mixed with pelleted cells for a final concentration of 3 – 5 x 10^6^ cells ml^-1^.

### 4.3 Hydrogel casting

Glass slides were hydrophobized with Sigmacote (SL2, Sigma Aldrich), washed with UPW, and sterilized by UV irradiation (λ=265 nm) in a biosafety cabinet. Casting molds (ø 6 mm x 2 mm high) were prepared from silicone mats (McMaster-Carr, Elmhurst, IL, USA) and sterilized by autoclavation. The silicone molds were firmly placed on the hydrophobized glass slides and resin was pipetted into the molds (55 µl per mold), covered by another hydrophobized glass slide and crosslinked under 20 mW cm^-2^UV irradiation (λ=365 nm, Thorlabs, Germany) for 5 min per side. After photocrosslinking, constructs were washed in prewarmed D-PBS and prepared for analyses or cell culture.

### 4.4 Volumetric bioprinting

Briefly, resin was transferred into the cylindrical ø 10 mm printing glass vials and chilled to 4 °C to form thermal crosslinking of GelMA allowing for stable printing. VBP of the different constructs was achieved using a Tomolite v1.0 equipped with a λ=405 nm laser in combination with Apparite software^[26]^ (both from Readily 3D, Lausanne, Switzerland). Glass vials were warmed to 37 °C after printing to remove unpolymerized resin and washed in prewarmed D-PBS. Then, constructs were transferred to 0.1% LAP in D-PBS and post-cured for 5 min under 20 mW cm^-2^UV irradiation (λ=365 nm, Thorlabs, Germany) and again washed in D-PBS before preparing them for analyses or cell culture. For prints with hMSCs, the process was done using sterile glass vials and reagents, and constructs were handled within a biosafety cabinet. Details on hMSC expansion and construct culture are given in the supplementary information. The branch model was provided by the printer manufacturer, the trabecular model was kindly provided by Dr. Riccardo Levato (Utrecht University), and the star model was freely available online (Prusa Research).

### 4.5 Analyses

Experimental details on the analyses can be found in the supplementary information.

### 4.6 Statistical analyses

Statistical analyses were performed, and graphs were prepared in GraphPad Prism (version 10.3.1, GraphPad, La Jolla, CA, USA). Data were tested for normality in distributions and equal variances using Shapiro-Wilk tests and Brown-Forsythe tests, respectively. When these assumptions were met, mean ± standard deviation are presented, and to test for differences, a one-way ANOVA (for the comparison >2 groups) followed by Holm-Šídák’s post hoc tests, or a two-way ANOVA (for comparisons between groups over time) followed by Tukey’s post hoc tests were performed. Other data are presented as median ± interquartile range and were tested for differences with non-parametric Mann–Whitney U tests or Kruskal-Wallis tests with Dunn’s post hoc tests with adjusted p-value for multiple comparisons. With a p-value of <0.05, differences were considered statistically significant.

## 6. Author contributions

Conceptualization: BdW, XHQ. Data curation: BdW. Formal analysis: BdW. Funding acquisition: BdW, XHQ. Resources: XHQ, RM. Investigation: BdW. Methodology: BdW, XHQ, MB, DZ. Project administration: BdW, XHQ. Supervision: XHQ, RM. Visualization: BdW. Writing – original draft: BdW. Writing – review & editing: BdW, XHQ, DZ, MB, RM.

## 7. Acknowledgement

This work was supported by a Rubicon Fellowship (BdW, no. 019.223EN.023) from the Dutch Research Council (NWO) as well as the NRP79 ‘‘Advancing 3R’’ grant of the Swiss National Science Foundation (XHQ, no. 206501). The authors thank Christian Gehre for methodological support, Jakob Dietz and Wanwan Qiu for experimental support. We gratefully acknowledge staff members of the ScopeM for technical support and assistance in this work. We thank Paul Delrot (Readily 3D) for his generous technical support and input on the scattering analysis, Riccardo Levato for providing the .STL file and Marc Bohner for scientific discussion.

## 8. Data availability statement

The data supporting the findings of this study will be made available in ETH Repository (DOI: 10.3929/ethz-b-000724832).

