## Supplementary Information for "Volumetric Bioprinting of Bone-like Mineralizing Hydrogel Constructs in the Presence of High Cell Densities and Mineral Precursors"

**Table of Content**

- Supplementary Table 1
- Supplementary Figures 1-17
- Supplementary Methods
- Supplementary References

### Supplementary Table

**Table S1.** Different resins used in this study.

| Resin mix | GelMA (%) | pAsp (mg $ml^{-1}$) | Iodixanol (% w/v) | $CaCl_2$ (mM) | LAP (%) | $K_2HPO_4$ (mM) | PBS | HEPES (mM) |
|---|---|---|---|---|---|---|---|---|
| GelMA | 5 | | | | 0.1 | | 1x | |
| GelMA + L pAsp | 5 | 0.1 | | | 0.1 | | 1x | |
| GelMA + H pAsp | 5 | 1 | | | 0.1 | | 1x | |
| GelMA + L salts | 5 | | | 9 | 0.1 | 4.2 | | 25 |
| GelMA + L salts + L pAsp | 5 | 0.1 | | 9 | 0.1 | 4.2 | | 25 |
| GelMA + L salts + H pAsp (GelMA + ACP) | 5 | 1 | | 9 | 0.1 | 4.2 | | 25 |
| GelMA + M salts | 5 | | | 18 | 0.1 | 8.4 | | 25 |
| GelMA + M salts + L pAsp | 5 | 0.1 | | 18 | 0.1 | 8.4 | | 25 |
| GelMA + M salts + H pAsp | 5 | 1 | | 18 | 0.1 | 8.4 | | 25 |
| GelMA + H salts | 5 | | | 27 | 0.1 | 12.6 | | 25 |
| GelMA + H salts + L pAsp | 5 | 0.1 | | 27 | 0.1 | 12.6 | | 25 |
| GelMA + H salts + H pAsp | 5 | 1 | | 27 | 0.1 | 12.6 | | 25 |
| GelMA + 10% iodixanol | 5 | | 10 | | 0.1 | | 1x | |
| GelMA + 15% iodixanol | 5 | | 15 | | 0.1 | | 1x | |
| GelMA + 20% iodixanol | 5 | | 20 | | 0.1 | | 1x | |
| GelMA + ACP + 10% iodixanol | 5 | 1 | 10 | 9 | 0.1 | 4.2 | | 25 |
| GelMA + ACP + 15% iodixanol | 5 | 1 | 15 | 9 | 0.1 | 4.2 | | 25 |
| GelMA + ACP + 20% iodixanol | 5 | 1 | 20 | 9 | 0.1 | 4.2 | | 25 |

Abbreviations: poly-aspartic acid (pAsp), gelatin methacryloyl (GelMA), low (L), medium (M), high (H), Lithium-Phenyl-2,4,6-trimethylbenzoylphosphinat (LAP), amorphous calcium phosphate (ACP).

**Supplementary Figures**

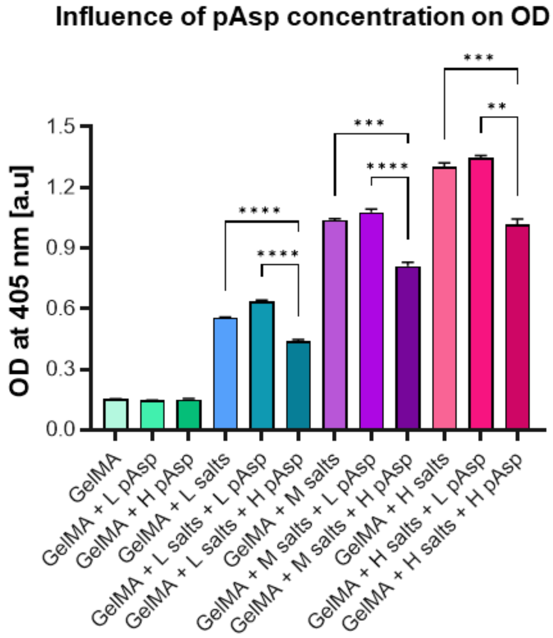


**Figure S1.** Different resins' optical density (OD) at the printer's wavelength ($\lambda$ = 405 nm) under influence of different salt (9 mM (L), 18 mM (M), or 27 mM (H) $CaCl_2$ and 4.2 mM (L), 8.4 mM (M), or 12.6 mM (H) $K_2HPO_4$) and pAsp (100 µg $ml^{-1}$ (L) or 1 mg $ml^{-1}$ (H)) concentrations, $N$ = 4, $p<0.05$ (Two-way ANOVA and Tukey's post hoc tests for comparisons between pAsp concentrations within the different salt conditions).

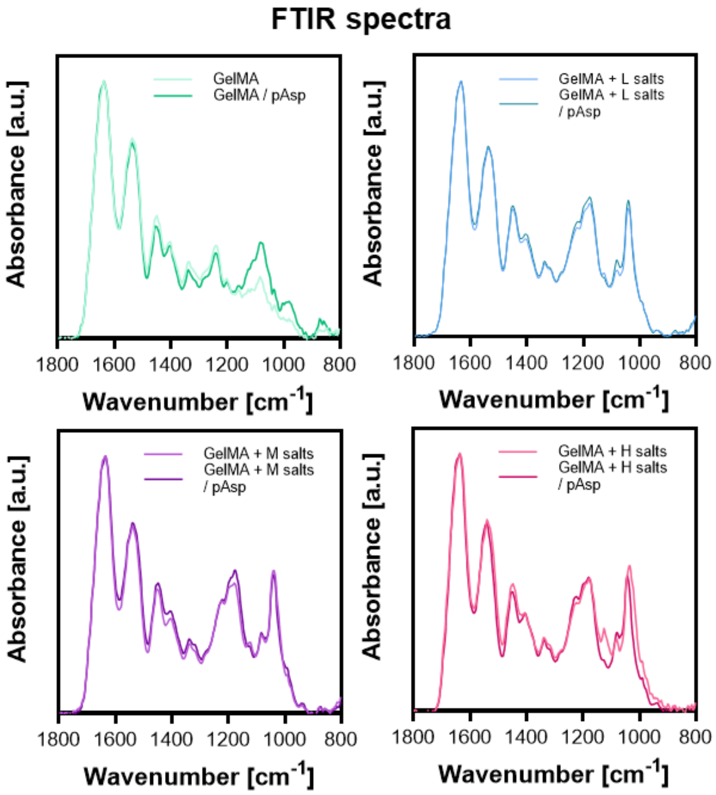


**Figure S2.** FTIR spectra per salt concentration and in the presence and absence of pAsp, $N$ = 3. Abbreviations: Fourier-transform infrared spectroscopy (FTIR), poly-aspartic acid (pAsp), gelatin methacryloyl (GelMA), low (L), medium (M), high (H).

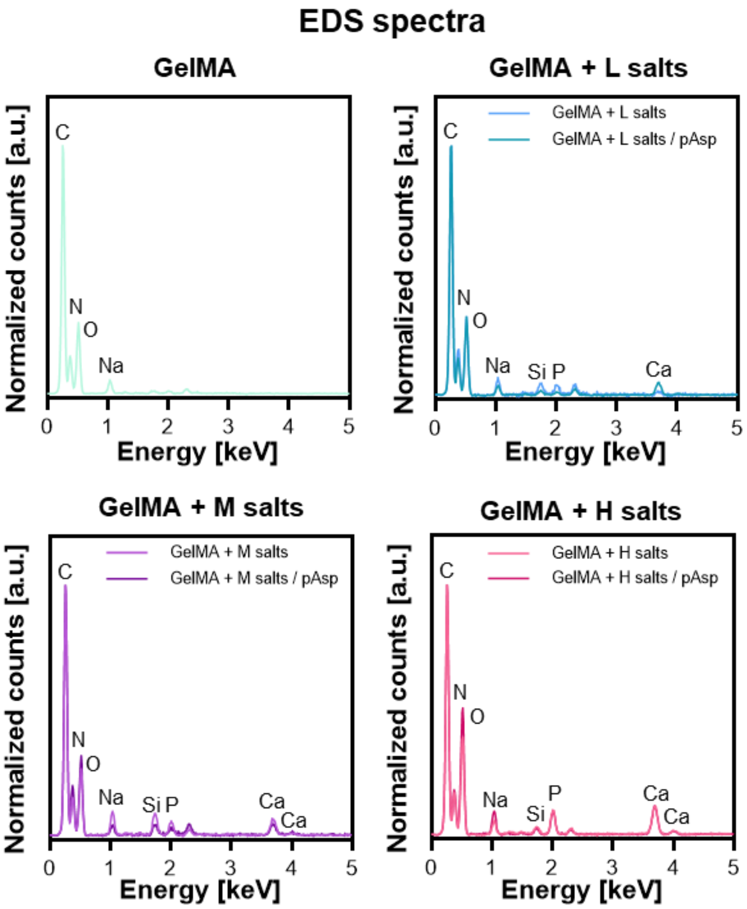


**Figure S3.** EDS spectra of hydrogel surfaces demonstrating the presence of calcium and phosphorus in all salt conditions, *N* = 3. Abbreviations: Energy dispersive spectroscopy (EDS), poly-aspartic acid (pAsp), gelatin methacryloyl (GelMA), low (L), medium (M), high (H).

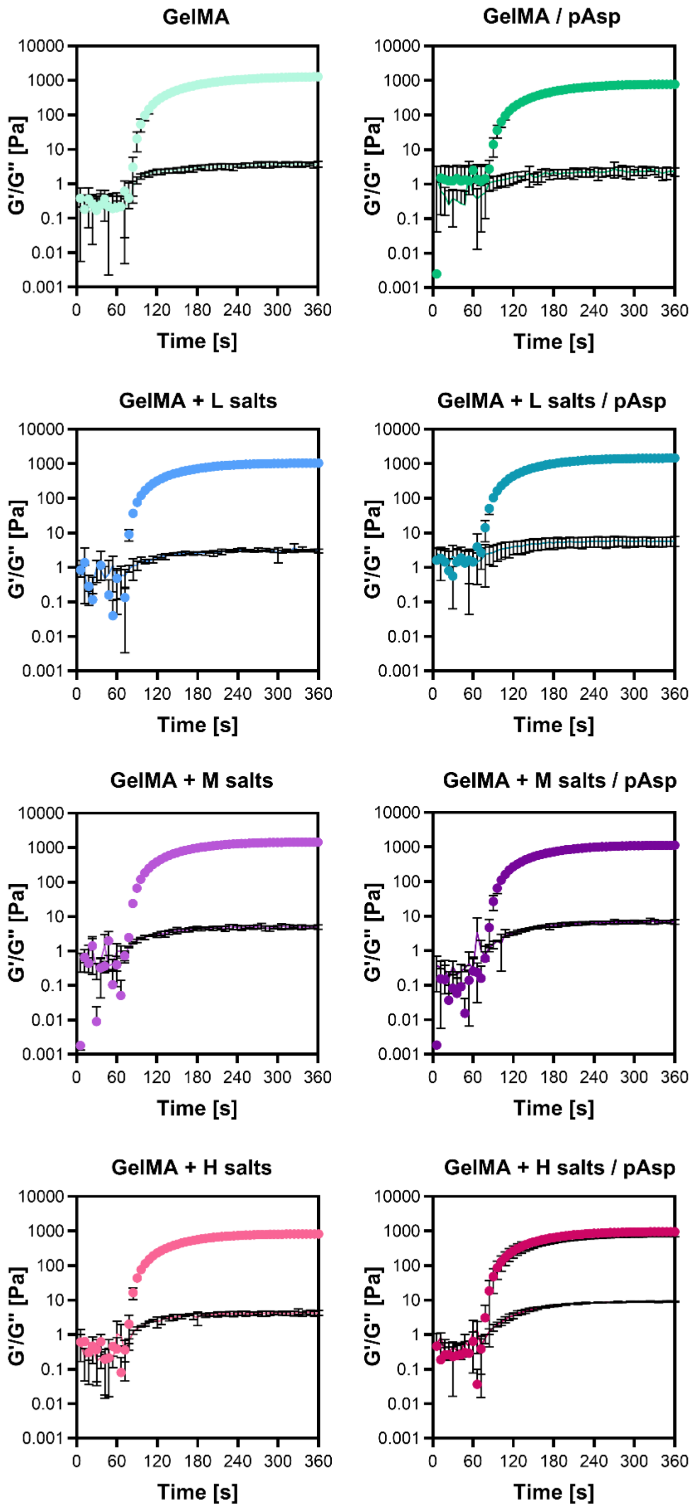


**Figure S4.** Storage and loss modulus measured under influence of UV-light (λ = 365 nm) from 60 s onwards, $N$ = 3. Abbreviations: poly-aspartic acid (pAsp), gelatin methacryloyl (GelMA), low (L), medium (M), high (H).

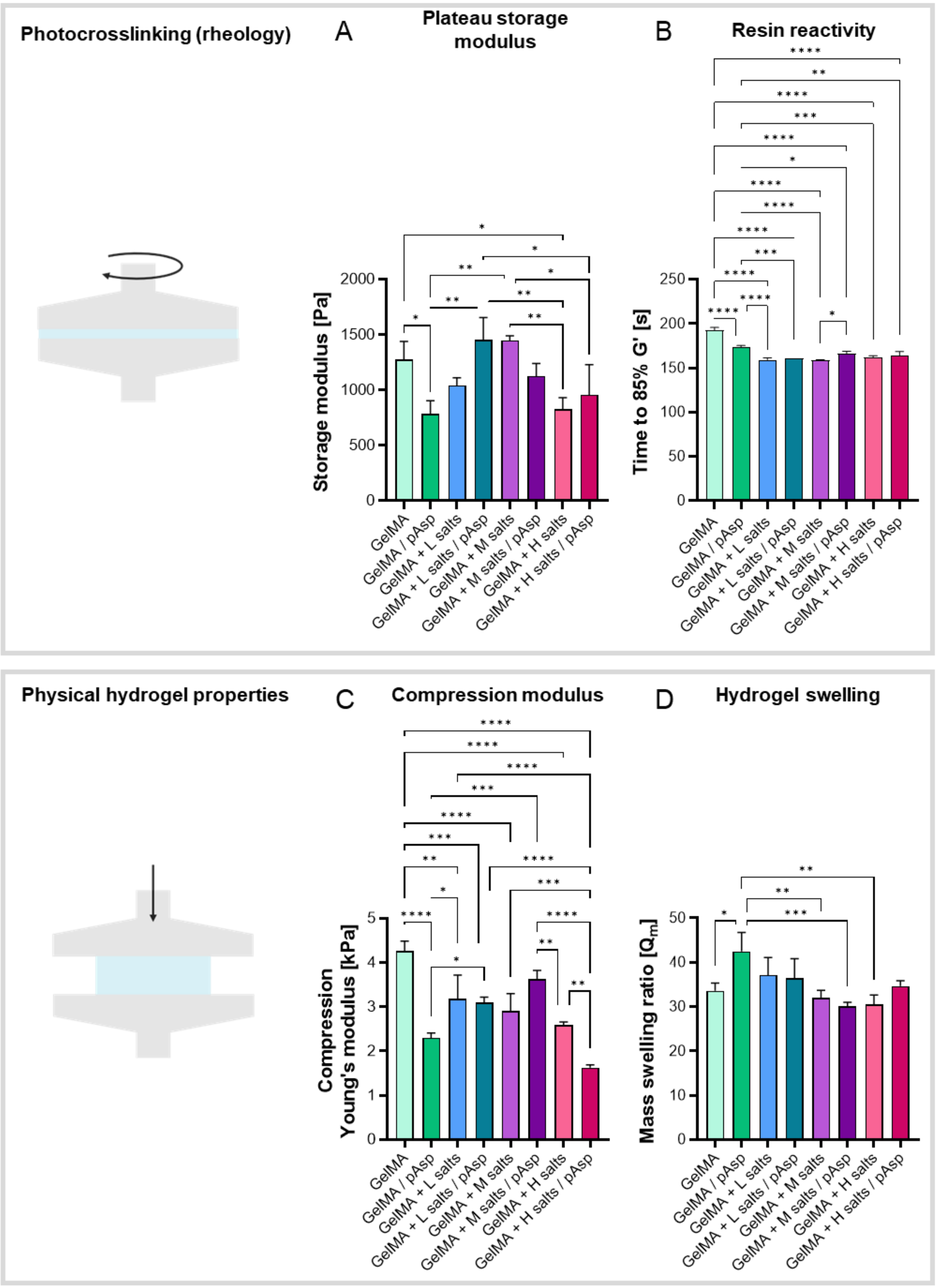


**Figure S5.** Crosslinking and physical properties of cast hydrogels. (**A**) Photo-rheology derived plateau storage modulus (G') (after 5 min photocrosslinking, *N* = 3, $p<0.05$ (One-way ANOVA and Holm-Šídák's post hoc tests). (**B**) Photo-rheology derived time to reach 85% of plateau storage modulus, *N* = 3, $p<0.05$ (One-way ANOVA and Holm-Šídák's post hoc tests). (**C**) Young's modulus derived from compression stress-strain curves, $N \geq 3$, $p<0.05$ (One-way ANOVA and Holm-Šídák's post hoc tests). (**D**) Cast hydrogel mass swelling ratio, $N \geq 3$, $p<0.05$ (One-way ANOVA and Holm-Šídák's post hoc tests). Abbreviations: poly-aspartic acid (pAsp), gelatin methacryloyl (GelMA), low (L), medium (M), high (H).

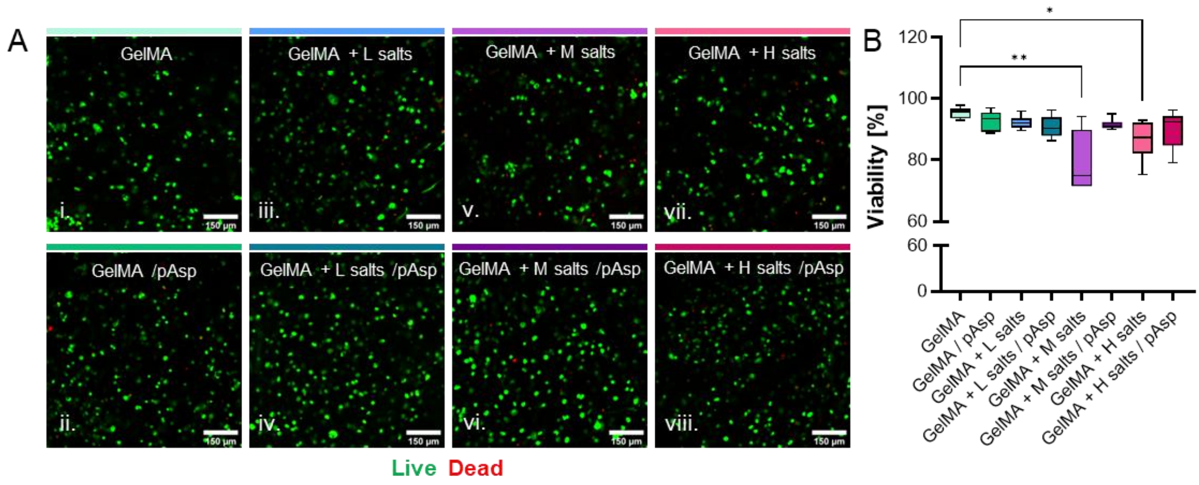


**Figure S6.** (**A**) Cell viability after 48 h of hMSC cultures (3 million cells $ml^{-1}$) embedded in cast hydrogels (i - viii), and the (**B**) quantification of viable cell percentages, $N$ = 3, $p$<0.05 (Kruskal-Wallis test and Dunn's post hoc tests). Asterisks in figures represent results of post hoc analyses (*$p$<0.05, **$p$<0.01, ***$p$<0.001, ****$p$<0.0001). Abbreviations: poly-aspartic acid (pAsp), gelatin methacryloyl (GelMA), low (L), medium (M), high (H).

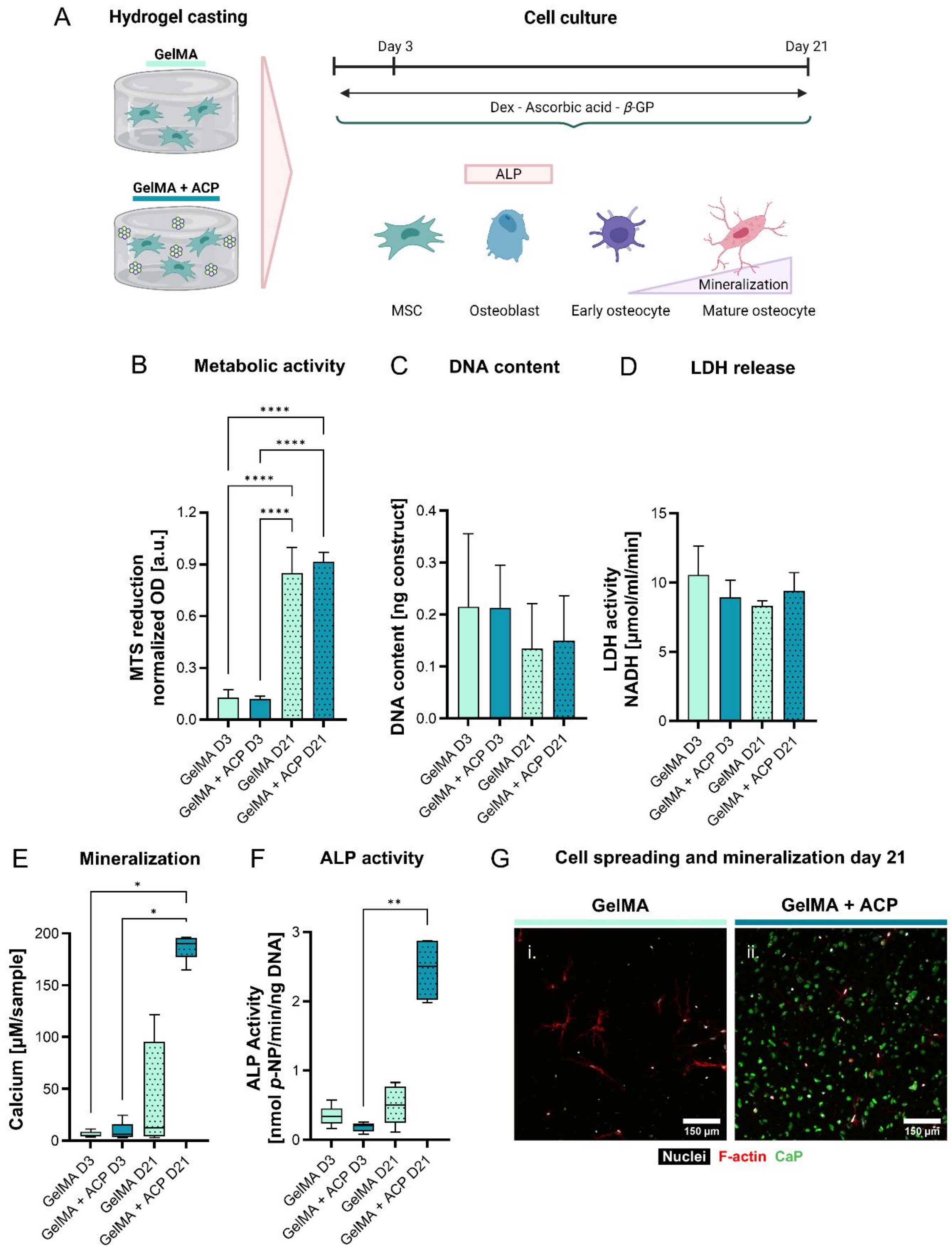


**Figure S7.** The influence of the cast resin formulations on osteogenic differentiation of hMSCs. (**A**) Constructs were cast and cultured for up to 21 days with osteogenic medium. Figure created with BioRender.com. (**B**) Cell metabolic activity measured by MTS reduction, $N \geq 4$, $p<0.05$ (One-way ANOVA and Holm-Šídák's post hoc tests). (**C**) Construct DNA content, $N \geq 4$, *ns* (One-way ANOVA and Holm-Šídák's post hoc tests). (**D**) LDH release in the medium as a measure for cell death, $N \geq 4$, *ns* (One-way ANOVA and Holm-Šídák's post hoc tests). (**E**) Construct mineralization measured by calcium content, $N \geq 4$, $p<0.05$

(Kruskal-Wallis with Dunn's post hoc tests). (**E**) Cellular ALP activity, $N \geq 4$, *ns* $p<0.05$ (Kruskal-Wallis with Dunn's post hoc tests). (**F**) Visualization of cell spreading and mineralization using CLSM after staining for nuclei, F-actin, and calcium phosphate, $N = 3$. Asterisks in figures represent results of post hoc analyses *$p<0.05$, **$p<0.01$, ****$p<0.0001$). Abbreviations: gelatin methacryloyl (GelMA), amorphous calcium phosphate (ACP), human mesenchymal stromal cells (hMSCs), lactate dehydrogenase (LDH), alkaline phosphatase (ALP), calcium phosphate (CaP).

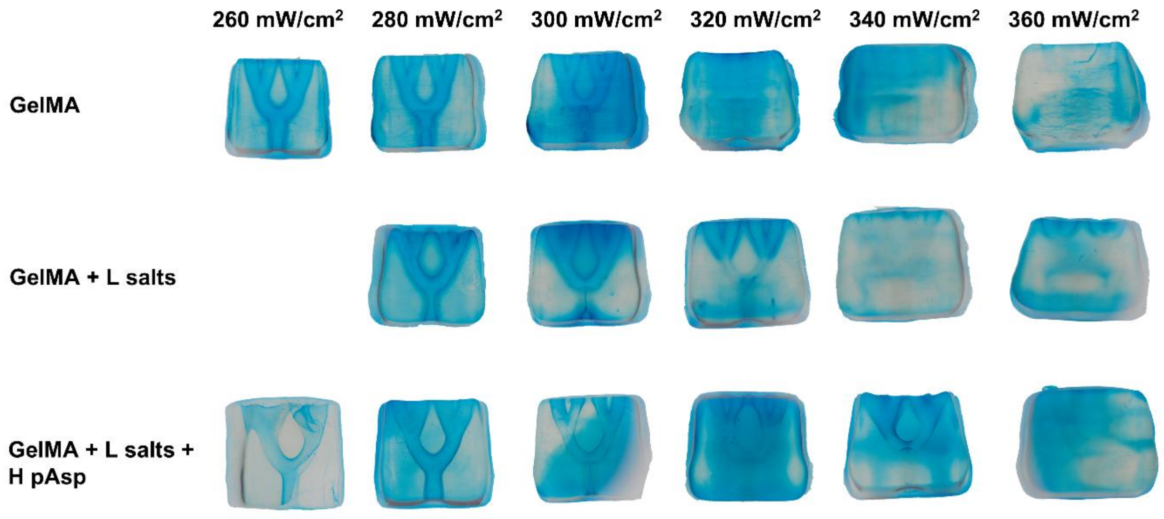


**Figure S8.** Printed branch constructs used for quantification of printability presented in **Figure 4G**. Abbreviations: poly-aspartic acid (pAsp), gelatin methacryloyl (GelMA), low (L).

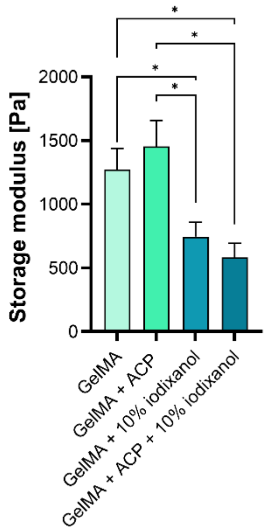


**Figure S9.** Photo-rheology derived plateau storage modulus (after 5 min photo-polymerization) for resins with 10% iodixanol and without, $N = 3$, $p<0.05$ (One-way ANOVA and Holm-Šídák's post hoc tests). Abbreviations: gelatin methacryloyl (GelMA), amorphous calcium phosphate (ACP).

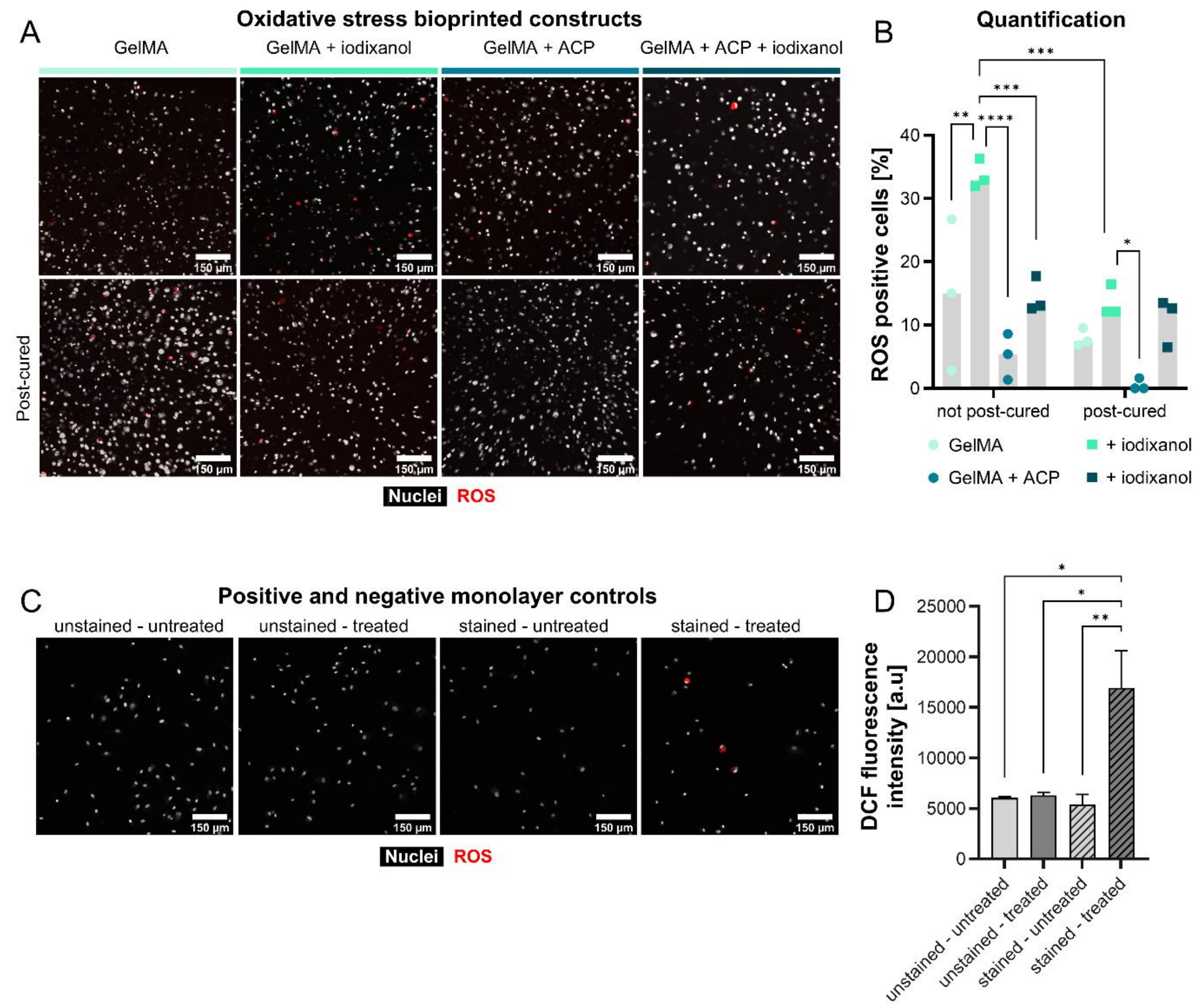


**Figure S10.** ROS and free radical experiments to study the role of iodixanol and post-curing on cellular oxidative stress. (**A**) Bioprinted constructs with DCFH-DA labelled hMSCs, post-cured and not post-cured stained for nuclei (Hoechst) and visualized with CLSM on the day of bioprinting. (**B**) Quantification of DCF positive cells, positive cells have been exposed to free radicals and/or ROS, *N* = 3, $p<0.05$ (Two-way ANOVA and Tukey's post hoc tests). (**C**) Positive and negative controls for ROS experiments that were (**D**) quantified by measuring the fluorescent intensity using a plate reader, *N* = 5, $p<0.05$ (One-way ANOVA and Holm-Šídák's post hoc tests). Abbreviations: gelatin methacryloyl (GelMA), amorphous calcium phosphate (ACP), reactive oxygen species (ROS).

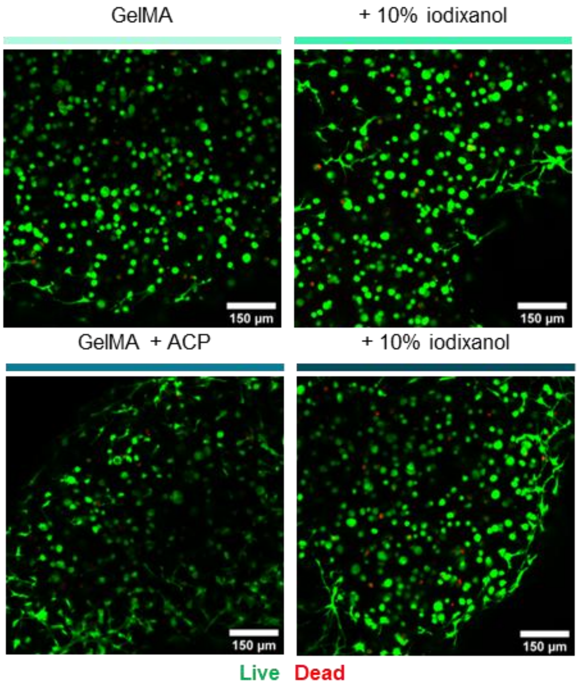


**Figure S11.** Cell viability of hMSCs (3 million cells $ml^{-1}$) 24 h post VBP. Abbreviations: human mesenchymal stromal cells (hMSCs), volumetric bioprintingin (VBP), gelatin methacryloyl (GelMA), amorphous calcium phosphate (ACP).

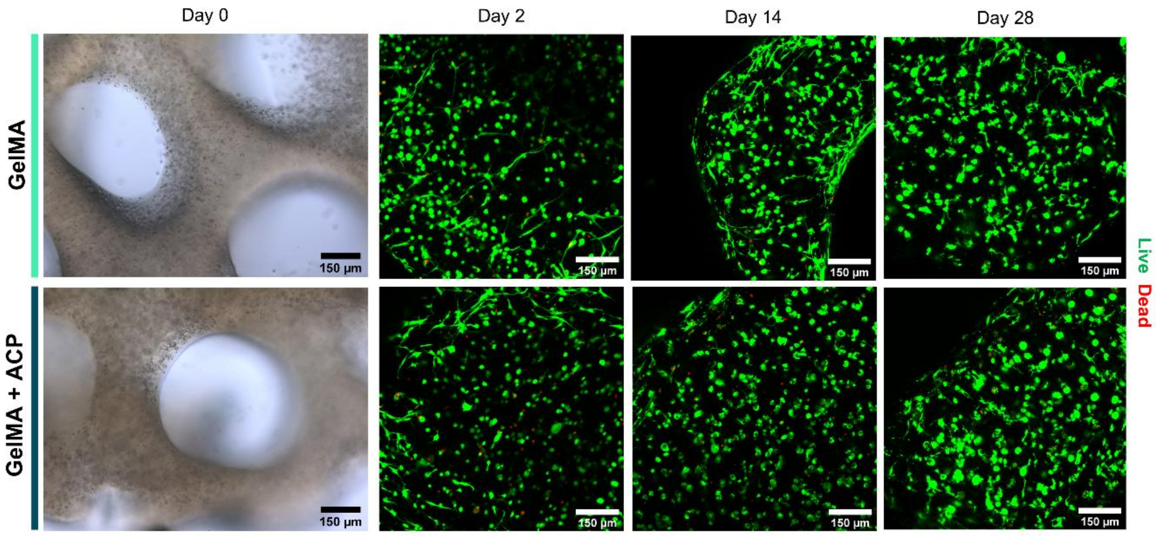


**Figure S12.** Bioprinted constructs and their cell viability over 28 days. Abbreviations: gelatin methacryloyl (GelMA), amorphous calcium phosphate (ACP).

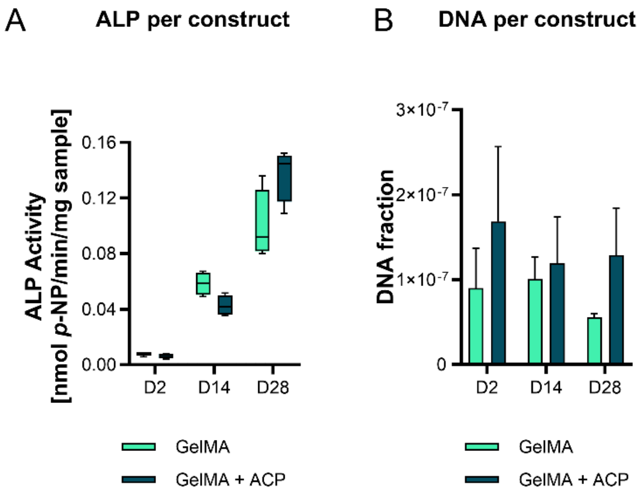


**Figure S13.** ALP activity and DNA content per construct. In the manuscript, the ALP content (**A**), has been corrected for DNA content per construct (**B**), *N* = 4. Abbreviations: alkaline phosphatase (ALP), gelatin methacryloyl (GelMA), amorphous calcium phosphate (ACP).

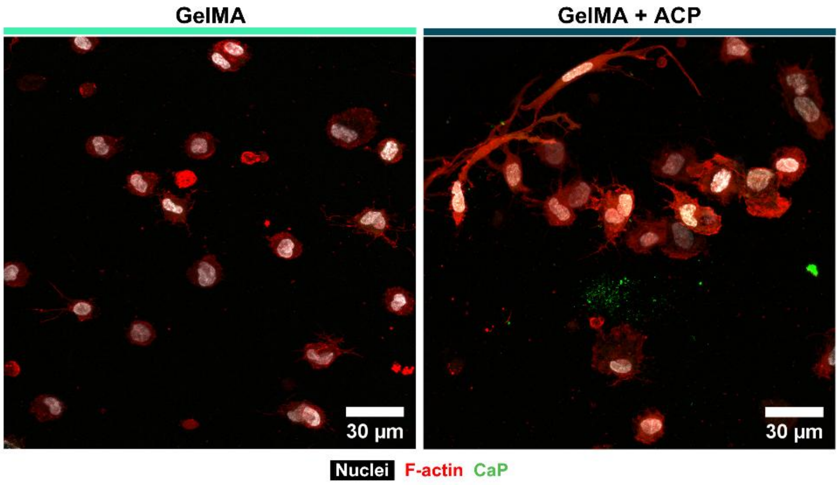


**Figure S14.** Visualization of cell spreading and mineralization in bioprinted constructs on day 2 using CLSM after staining for nuclei, F-actin, and calcium phosphate, *N* = 3. Abbreviations: gelatin methacryloyl (GelMA), amorphous calcium phosphate (ACP), calcium phosphate (CaP).

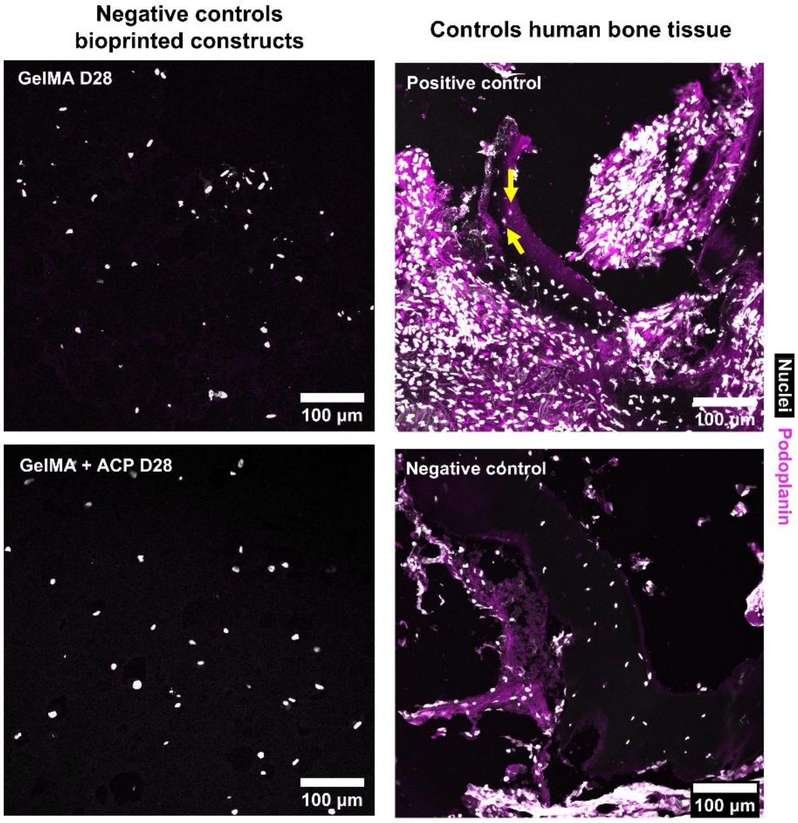


**Figure S15.** Positive (human bone) and negative (only secondary antibody) controls for immunohistochemistry of podoplanin. Podoplanin was visible in the positive control in the osteocyte cell processes (yellow arrows). Abbreviations: day (D), gelatin methacryloyl (GelMA), amorphous calcium phosphate (ACP).

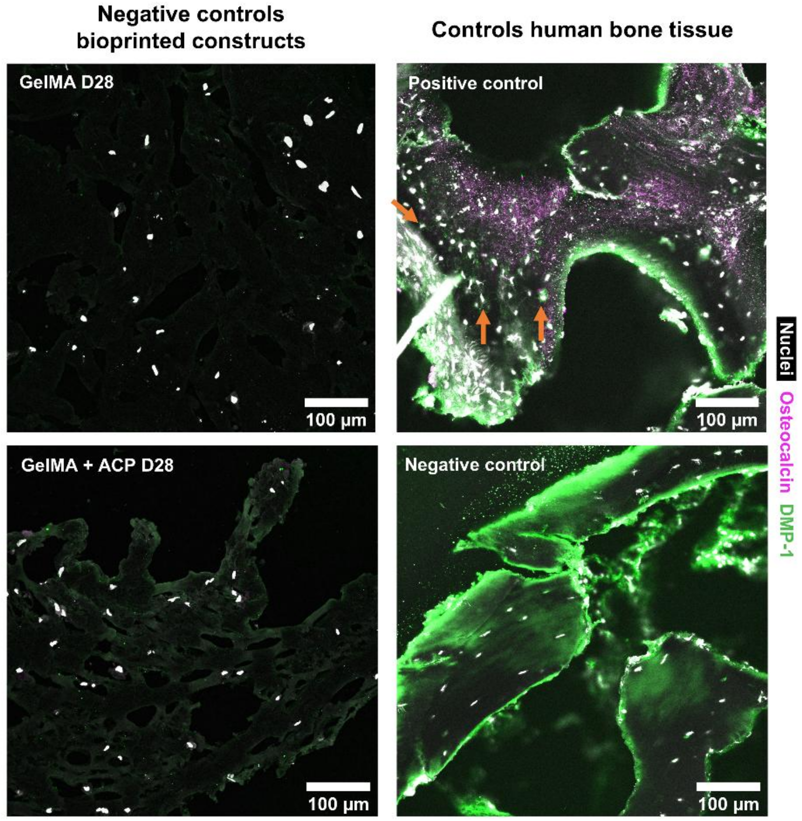


**Figure S16.** Positive (human bone) and negative (only secondary antibody) controls for immunohistochemistry of osteocalcin and DMP1. Osteocalcin was visible in the positive controls in the matrix, DMP1 was visible in the positive control in the osteocyte cell processes (orange arrows). Abbreviations: day (D), gelatin methacryloyl (GelMA), amorphous calcium phosphate (ACP), dentin matrix acidic phosphoprotein 1 (DMP1).

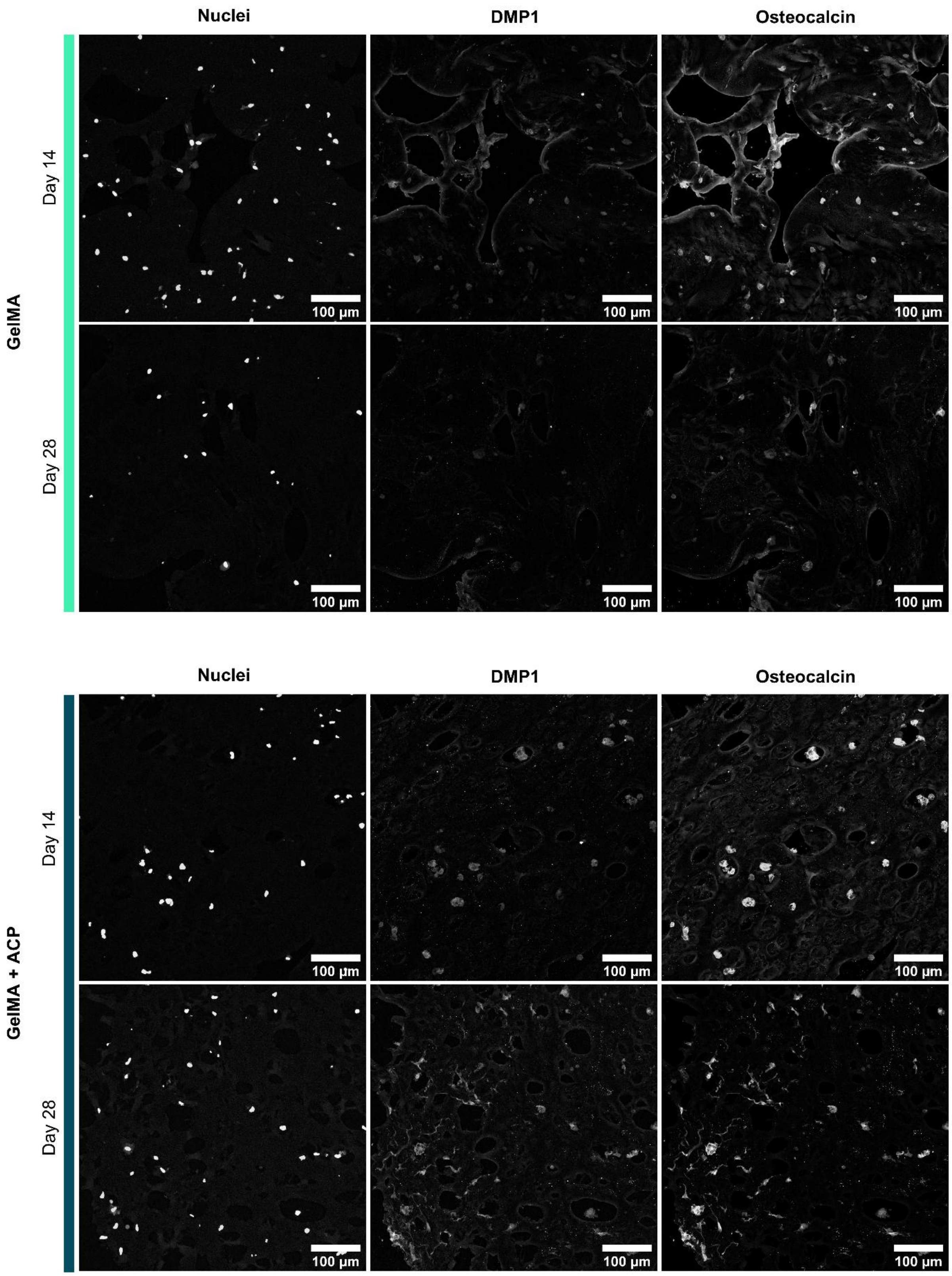


**Figure S17.** Single channel results for DMP1 and osteocalcin antibody stainings. Abbreviations: gelatin methacryloyl (GelMA), amorphous calcium phosphate (ACP), dentin matrix acidic phosphoprotein 1 (DMP1).

### Supplementary Methods

#### *Mesenchymal stromal cell expansion and construct culture*

For cell expansion, hMSCs (Male donor, PT-2501, Lonza, Walkersville, MD, USA) were seeded at a density of 2.5*$10^3$ cells $cm^{-2}$ and cultured in expansion medium containing high glucose Dulbecco's modified Eagle's medium (hg-DMEM; 41966029, Thermo Fisher Scientific), 10% FBS (10270106, lot# 2440094, Thermo Fisher Scientific), 1% anti-anti (15240062, Thermo Fisher Scientific), 1% Non-Essential Amino Acids (11140, Thermo Thermo Fisher Scientific), and 1 ng/mL basic fibroblastic growth factor (13256-029, Thermo Fisher Scientific) at 37 ºC and 5% $CO_2$. At 80% confluency, cells were detached using 0.25% trypsin-EDTA (25200-056, Thermo Fisher Scientific) and were used for experiments at passage 4-7. For culturing cast or bioprinted hydrogels, constructs were cultured in osteogenic medium containing low glucose (lg)-DMEM (11880036, Thermo Fisher Scientific), 10% FBS, 1% anti-anti, 2 mM glutamax (35050038, Thermo Fisher Scientific), 25 mM HEPES supplemented with 50 µg $mL^{-1}$ ascorbic acid-2-phosphate (A8960, Sigma Aldrich), 100 nM dexamethasone (D2915 Sigma Aldrich), and when indicated with 10 mM β-glycerophosphate (410991000, Thermo Fisher Scientific). Medium was changed three times a week.

#### *OD measurements*

The OD was measured as the absorption at $\lambda$=405 nm with a plate reader (Spark M10, Tecan). Per well, 100 µl of resin was cast and dependent of the measurement, *i.e.,* single or over a period, in the presence or absence of LAP, respectively. For single measurements, resins were cooled to 4 °C to match the printing temperature.

#### *SEM, BSE detection and EDS*

Samples were fixed in 2.5% glutaraldehyde (G5882, Sigma Aldrich) in 0.1 M sodium cacodylate buffer (CB) for 30 min and then washed in CB. Samples were further processed using a BioWave Pro+ Tissue Processor (Pelco) and additionally washed thrice in CB, in case of cellular samples stained with 1% osmium tetroxide solution (251755, Sigma Aldrich) in double distilled $H_2O$ (dd$H_2O$) under vacuum, washed thrice with dd$H_2O$, and dehydrated in graded ethanol series. For acellular samples, osmium staining was omitted to avoid interference during EDS measurements. Samples were dried using a CPD 931 Critical Point Dryer (Tousimis), mounted on subs with carbon tape, grounded with Ag-paint, and coated with a 10 nm layer carbon (CCU-010 Carbon Coater, Safematic). SEM and BSE imaging were performed in high vacuum with a 10 kV electron beam (Quanta 200F, Thermo Fisher Scientific). EDS spectra and maps were obtained (EDAX Octane Super) using a 20 kV electron beam. Upon spectrum collection, baselines were subtracted using Origin 2021 (OriginLab Corp., Northampton, MA, USA) and data was normalized to the carbon peak.

***FTIR***

Cast hydrogels were washed thrice in UPW, frozen at -80 °C, and lyophilized to dehydrate the samples. FTIR spectra were collected from 800 $cm^{-1}$ to 1800 $cm^{-1}$ by averaging 32 spectra with a resolution of 4 $cm^{-1}$ (Varian 640). The background was measured and subtracted from the spectra. Upon spectrum collection, baselines were subtracted using Origin 2021 and spectra were normalized to the amide I peaks.

***Photo-rheology***

Resins were pipetted (70 µl) on the bottom glass plate on a rheometer (MCR 302, Anton Paar, Graz, Austria), equipped with a sand-blasted PP20 measuring plate and a UV-lamp (λ=365 nm) for photocrosslinking at 20 mW $cm^{-2}$. To prevent dehydration during the measurements, the plate was surrounded by mineral oil (330779, Sigma Aldrich). Measurements were done at a working distance of 0.1 mm and ambient temperature of 25 °C. Storage and loss moduli were measured during polymerization at 1 Hz oscillations with 0.5% strain. After 60 s, UV-light was turned on and the measurements continued for 5 more min. The final storage modulus values were used as storage modulus plateau value, 85% of this value was calculated and the time it took to reach this value was used to provide a measurement for resin reactivity.

***Compression test***

Cast hydrogels (ø 6 mm x 2 mm high) were tested in unconfined compression at a Zwick mechanical test system (Zwick, 1456, Ulm, Germany) equipped with a 10 N load cell. Samples were preloaded at 5 mN and then subjected to 10% strain $min^{-1}$ up to 30 % strain. The compression Young's modulus was determined by the slope of a linear fit to the load-displacement curves between 2% and 10% displacement.

***Swelling test***

The mass swelling ratio ($Q_m$) of the hydrogels was determined by measuring the wet weight ($M_w$) after 48 hours of swelling in UPW and dividing this by their dry weight ($M_d$), which was obtained by after lyophilization (**Equation 1**).

$$Q_m = \frac{M_w}{M_d} \quad \text{Equation 1}$$

***Resin printer light transmission and scattering visualization***

Light transmission through the resins was measured by quantifying the light power loss within the Tomolite bioprinter using an optical power meter (Thorlabs). Cuvettes were filled with UPW or resin and the light transmitted through the resin was collected for 40 s, with the spatially coherent light beam ($\lambda$ = 405 nm) switched on from 10 s. Scattering within resins was visualized by projecting a spatially coherent light beam on resin-filled cylindrical printing vials (ø 10 mm) and by taking photographs with the orthogonal camera within the printer.

***Print evaluation***

Printed constructs were visually evaluated by staining (stars and trabeculae) or perfusing (branch models) with alcian blue solution (1.01647, Sigma Aldrich) and by imaging them with a stereo microscope (Olympus, Japan).

***Refractive index measurements***

Refractive index measurements were performed by loading 300 µl resin onto a refractometer (ORL 94BS, Kern, Stuttgart, Germany) after calibration with UPW at ambient conditions.

***Cellular ROS assay***

The cellular ROS was evaluated using 2′,7′-Dichlorodihydrofluorescein diacetate (DCFH-DA) labeling following a protocol modified from literature.[1] DCFH-DA was dissolved in DMSO at a 50 mM concentration to prepare a 500x stock concentration. Single use stock-aliquots were stored at -20 °C. Prior to labeling, cells were washed in PBS and resuspended in 2 ml per 5 million cells labeling solution containing 100 µM DCFH-DA in serum and phenol red free DMEM (11880036, Thermo Fisher Scientific). Cells were labelled in a shaking water bath at 37 °C for 40 min. After labelling, cells were centrifuged to remove the labelling solution and cells were washed thrice in PBS. Cells were then processed for VBP as described in Sections 5.2 and 5.4. To then evaluate the printing and post-curing induced oxidative stress, cells were counter-stained with Hoechst (1:500, B2261, Sigma Aldrich) and imaged using CLSM (Zeiss LSM 780 with 10/0.3 EC Plan-Neofluor objective) using the FS49 (Hoechst, $\lambda_{ex}$ = 365 nm, $\lambda_{em}$ = 445/50 nm) and FS38 (DCF, $\lambda_{ex}$ = 470 nm, $\lambda_{em}$ = 525/50 nm) filters. Maximum intensity projections were created and analysed in FiJi.[2] The total cell number was evaluated using particle analysis while the number of ROS positive cells was manually counted. The percentage of ROS positive cells was calculated as the fraction of ROS positive cells over the total cell number. As control experiments, labeled and unlabeled cells were seeded in clear (microscopy) and black (plate reader) 96-well plates (10,000 cells well$^{-1}$), and left untreated or treated with 500 µM $H_2O_2$ for 1 h. The clear assay plates were counter-stained and imaged with CLSM as described above. The black assay plates were evaluated for their fluorescence intensity using a plate reader (Spark M10, Tecan) at λex = 485/20 nm, λem = 535/25 nm.

***Cell viability***

To assess cell viability, constructs were washed in PBS and subsequently stained with 2 µM Calcein Green AM (56496, Sigma Aldrich) and 4 µM Ethidium-homodimer-1 (460439, Sigma Aldrich) in phenol red free DMEM (11880036, Thermo Fisher Scientific) for 20 min at 37 °C. After staining, constructs were washed in PBS and z-stacks were collected in phenol red free DMEM using CLSM (Zeiss LSM 780 with 10/0.3 EC Plan-Neofluor objective). Maximum intensity projections were created and analysed in FiJi.[2] Cell viability was evaluated using particle analysis for both the Calcein-AM (live) and Ethidium-homodimer-1 (dead) channel. The percentage of viable cells was calculated as the fraction of live cells over the sum of live and dead cells.

***LDH assay***

A 100 *µ*l medium sample or NADH (10107735001, Sigma-Aldrich) standard was incubated with 100 *µ*l LDH reaction mixture (11644793001, Sigma-Aldrich) in 96-wells assay plates. Absorbance was measured after 5, 10 and 20 min at 490 nm (Spark M10, Tecan), and LDH activity was calculated between 10- and 20-min reaction, using standard curve absorbance values. Medium that has not been in contact with cells, was used as a control.

***MTS assay***

To evaluate cellular metabolic activity, MTS (3-(4,5-dimethylthiazol-2- yl)-5-(3-carboxymethoxyphenyl)-2-(4-sulfophenyl)-2H-tetrazolium) (ab197010, Abcam) reduction was measured. Constructs were incubated in 10% v/v MTS solution for 4 h at 37 °C and 5% $CO_2$ in the dark. Absorbance was read at 490 nm (Spark M10, Tecan). Medium that has not been in contact with cells, was used as a control and these values were subtracted from cellular samples.

***Calcium assay***

Hydrogels were homogenized using Fisherbrand™ Pellet Pestle™ and incubated in 0.5M acetic acid for 48 h at room temperature on a shaker. For stock solutions, a 100 µg ml$^{-1}$ O-cresolphthalein complexone (182440050, Thermo Fisher Scientific) solution was prepared in boric acid buffer (pH 8.5) and stored at 4 °C for up to two weeks, and a 1% w/v 8-hydroxyquinoline (252565, Sigma Aldrich) solution was prepared in 95% ethanol and stored at room temperature in the dark. To initiate the reaction after sample incubation, 10 µl supernatant was transferred to a 96-well assay plate and mixed with 300 µl freshly prepared reaction buffer containing 0.05 parts O-cresolphthalein complexone solution, 0.02 parts 8-hydroxyquinoline solution, and 0.05 parts 14.8 M ethanolamine (02400, Sigma Aldrich) in boric acid buffer (pH 11). After 10 minutes incubation at room temperature, absorbance was read at 570 nm and calcium content was quantified using standard curve absorbance values.

***ALP activity assay***

Hydrogels were homogenized using Fisherbrand™ Pellet Pestle™ and incubated in cell lysis buffer containing 0.2% (v/v) Triton X-100 and 5 mM $MgCl_2$. ALP activity in cell lysates was determined by adding 20 µl of 0.75 M 2-amino-2-methyl-1-propanol (A65182, Sigma-Aldrich) to 80 µl sample in 96-wells assay plates. Subsequently, 100 µl substrate solution (10 mM p-nitrophenyl-phosphate (71768, Sigma-Aldrich) in 0.75 M 2-amino-2-methyl-1-propanol) was added and wells were incubated at room temperature for 15 minutes. To stop the reaction, 100 µl 0.2 M NaOH was added. Absorbance was measured with a plate reader at 405 nm and these values were converted to ALP activity (converted p-nitrophenyl phosphate in µmol/ml/min) using standard curve absorbance values. ALP activity was normalized for DNA content.

***DNA assay***

After completing the ALP assay, remaining cell lysates underwent three cycles of freeze-thawing and ultrasonification. Then, samples were incubated at room temperature for 48 h. DNA content was quantified using a Quant-IT PicoGreen dsDNA assay (P7589, Thermo Fisher Scientific) according to the manufacturer's instructions. Briefly, 87.5 µl of 1x TE buffer, 12.5 µl of sample and 100 µl of PicoGreen working solution were added to black bottom 96-well assay plates. Fluorescence intensity was measured using a plate reader (Spark M10, Tecan) at $\lambda_{ex}$ = 485/20 nm, $\lambda_{em}$ = 535/25 nm. DNA content was derived from measuring fluorescence intensity for standard curve values.

***Immunofluorescent labeling***

Constructs were washed in PBS and fixed in 4% paraformaldehyde in PBS for 1 h and washed again in PBS.

For rhodamine and mineral labelling of bioprinted constructs, Methacryloxyethyl thiocarbamoyl rhodamine B (Polysciences, Hirschberg an der Bergstrasse, Germany) stock in DMSO (10 mg $ml^{-1}$) was dissolved at a concentration of 1 µg $ml^{-1}$ in 0.1% LAP in PBS. Printed constructs were placed in this labeling solution and incubated for 5 min under 20 mW $cm^{-2}$ UV irradiation (λ=365 nm, Thorlabs, Germany). After rhodamine labelling, samples were washed in PBS and minerals were stained using OsteoImage™ (PA-1503, Lonza) 1:100 for 30 min. Samples were subsequently washed and imaged in PBS using CLSM (Leica SP8, 10x/0.3 HC PL FLUOTAR objective).

For whole mount staining of nuclei, F-actin, and minerals, samples were permeabilized for 20 min in 0.5% v/v triton X-100 in PBS, followed by 2 h staining in 66 nM Alexa Fluor 647-conjugated Phalloidin (A30107, Thermo Fisher Scientific), Hoechst, and OsteoImage™. After staining, samples were washed and imaged in PBS using CLSM (Zeiss LSM 780 with 10/0.3 EC Plan-Neofluor and 40x/1.1 LD C-Apochromat objective).

For cryosections, samples were incubated in 5% (w/v) sucrose in PBS for 15 min followed by incubation in 35% (w/v) sucrose in PBS for 15 min and an incubation in a 1:1 mix of Tissue Tek® (Sakura) and the 35% sucrose solution for 15 min. Gels were then transferred to cryomolds, embedded in Tissue Tek® and frozen on a cooling dish floating on liquid $N_2$. Cryosections were sliced with a thickness of 30 µm and dried overnight and kept at -20 ºC until staining. Upon staining, sections were washed twice with PBS to remove Tissue Tek®. Then, sections were permeabilized in 0.3% v/v triton X-100 in PBS for 20 min. Non-specific antibody binding was then blocked with 1% w/v bovine serum albumin (BSA) and 5% v/v secondary antibody host-serum for 45 min (in case of multiple hosts, a 1:1 mixture was prepared). Primary antibodies (anti-osteocalcin: 1:200 (ab93876, Abcam, rabbit polyclonal), anti-DMP1: 1:100 (sc-73633, lot# I0122, Santa Cruz Biotechnology, mouse monoclonal $IgG_1$), and anti-podoplanin: 1:100 (sc-59347, lot# L0221, Santa Cruz Biotechnology, mouse monoclonal IgG)) were diluted in blocking buffer and sections were incubated overnight at 4 °C with primary antibody or blocking buffer (as negative control). After incubation, antibody solution was removed, and sections were washed in thrice in 0.025% v/v Triton X-100 in PBS (wash buffer). Sections were subsequently incubated with corresponding secondary antibodies 1:200 (DMP1: Donkey-anti mouse IgG Alexa Fluor 647 (ab150107, Abcam), osteocalcin: Goat-anti Rabbit IgG Alexa Fluor 555 (ab150082, Abcam), podoplanin: Donkey-anti mouse IgG Alexa Fluor 647 (ab150107, Abcam)) with Hoechst 1:500 in 1% BSA in 0.3% Triton X-100 in PBS for 2 h at room temperature. Sections were subsequently washed thrice in PBS and mounted with ProLong™ Gold Antifade Mountant (P10144, Thermo Fisher Scientific). Samples were imaged using CLSM (Leica SP8, 20x/0.7 HC PLAN APO objective). All images were prepared in FiJi.[2]
